# *De novo* design of parallel and antiparallel A_3_B_3_ heterohexameric α-helical barrels

**DOI:** 10.1101/2024.09.25.614870

**Authors:** Joel J. Chubb, Katherine I. Albanese, Alison Rodger, Derek N. Woolfson

**Affiliations:** School of Chemistry, University of Bristol, Cantock’s Close, Bristol BS8 1TS, UK; School of Natural Sciences, Macquarie University, Sydney, New South Wales, Australia; Research School of Chemistry, Australian National University, ACT, 2601, Australia; Max Planck-Bristol Centre for Minimal Biology, University of Bristol, Cantock’s Close, Bristol BS8 1TS, UK; School of Biochemistry, University of Bristol, Medical Sciences Building, University Walk, Bristol BS8 1TD, UK; Bristol BioDesign Institute, University of Bristol, Cantock’s Close, Bristol BS8 1TS, UK

**Keywords:** coiled coil, computational design and modelling, peptide design, peptide synthesis, self-assembly, synthetic biology

## Abstract

The *de novo* design of α-helical coiled-coil peptides is advanced. Using established sequence-to-structure relationships, it is possible to generate various coiled-coil assemblies with predictable numbers and orientations of helices. Here we target new assemblies, namely A_3_B_3_ heterohexamer α-helical barrels. These designs are based on pairs of sequences with 3-heptad repeats (***abcdefg***) programmed with ***a*** = Leu, ***d*** = Ile, ***e*** = Ala, and ***g*** = Ser, and ***b*** = ***c*** = Glu to make the acidic (A) chains and ***b*** = ***c*** = Lys in the basic (B) chains. These design rules ensure that the desired oligomeric state and stoichiometry are readily achieved. However, controlling the orientation of neighboring helices (parallel or anti-parallel) is less straightforward. Surprisingly, we find that assembly and helix orientation are sensitive to the starting position of the heptad repeats (the register) in the peptide sequences. Peptides starting at ***g*** (***g***-register) form a parallel 6-helix barrel in solution and in an X-ray crystal structure, whereas the ***b***- and ***c***-register peptides form an antiparallel complex. In lieu of experimental X-ray structures for ***b***- and ***c***-register peptides, AlphaFold-Multimer is used to predict atomistic models. However, considerably more sampling than the default value is required to match the models and the experimental data, as many confidently predicted and plausible models are generated with incorrect helix orientations. This work reveals the previously unknown influence of heptad register on the orientation of *α*-helical coiled-coil peptides and provides insights for the modeling of oligopeptide coiled-coil complexes with AlphaFold.

## INTRODUCTION

The *de novo* design of proteins and protein-like peptide assemblies has advanced significantly in recent years.^1-3^ Over the past two decades, we and others have aimed to understand and parameterize a particular type of peptide assembly and protein fold called the α-helical coiled coils (CCs).^4-6^ This has resulted in a guided exploration of CC sequence and structure space with *de novo* designed peptide and protein assemblies.^7-11^ From this, we have developed a set of rules to design peptides that assemble into a wide range of CC structures.

CCs comprise two or more α helices that wrap around one another like threads of rope creating a larger super helix.^12-14^ A key component of CC assembly is that hydrophobic side chains of one α helix project into diamond-shaped holes formed between side chains of a neighboring helix. These are called knob-into-hole (KIH) interactions and they specify CC structures in terms of oligomeric state, helix-partner preferences, and helix orientation.^4^ KIH interactions are encoded by sequence repeats of hydrophobic (***h***) and polar residues (***p***), (***hpphppp***), often called heptads and labelled ***abcdefg***. It is the ***h*** residues at ***a*** and ***d*** that form the main hydrophobic KIH interactions, but, as described below, residues at ***e*** and ***g*** also contribute to the helical interfaces through additional hydrophobic and salt-bridging interactions. Typically, stable CC peptide assemblies require 3 or more contiguous heptad repeats, and the canonical ***hpphppp*** repeats with charged residues at ***e*** and ***g*** tend to form dimers, trimers, or tetramers.

α-Helical barrels (αHBs) are a subset of CCs in which 5 or more α-helices assemble to generate central solvent-accessible channels along the super-helical axis.^8, 15^ Most αHBs are programmed by variations of the heptad repeat with further interfacial, ***h*-**like residues at the ***e*** and ***g*** sites to give ***hpphhph*** repeats. Over the last decade, we have determined clear sequence-to-structure relationships that define the oligomeric states for specific assemblies of 5 – 9 helices, which can be used as rules for rational and computational *de novo* peptide and protein design.^4, 8, 9, 11^ These rules are exemplified in the next section for hexameric barrels. The *de novo* αHBs presented to date are also highly thermostable making them exciting prospects as scaffolds for functional protein design. Indeed, they have been used for small-molecule sensing, ion transport across membranes, and rudimentary catalysis.^16-18^ Recently, single-chain variants of αHBs with 5 – 8 helices with pseudo-cyclic symmetry have been generated by using the sequence-to-structure relationships as seeds for AI-based computational design.^11^

Whilst heteromeric dimeric, trimeric and tetrameric CCs have been designed,^10, 19-24^ most of *de novo* designed αHB assemblies are homomeric. For functionalization and applications, heteromeric assemblies have advantages including controlled assembly—*i*.*e*., conditional folding dependent on the presence of 2 or more different peptide/protein chains—and a reduction in sequence and structural symmetry allowing subsets of the peptides to be modified. Mostly, heteromeric *de novo* CCs have been targeted by using different charged variants of the peptide chains to promote discriminating electrostatic interactions between the different chains. For instance, mixtures of acidic (A, anionic) and basic (B, cationic) peptides have been used to create AB-type dimers, ABC trimers, and A_2_B_2_ tetramers.^19-21^ Others have used cofactors, steric packing and hydrogen-bonding networks to generate heteromeric CCs.^24-27^ However, these require mutation of core-packing residues. There have been some designs of A_3_B_3_ αHBs and bundles, however, our previous attempt to do this gave homomeric assemblies of either the A or B peptides as well as the desired heteromeric barrels, which complicates the system for downstream applications (Figure 1).^28^ Similar behavior is observed for a heterohexameric bundle designed by Spencer and Hochbaum.^29, 30^ To our knowledge, the design of conditional heteromeric assemblies of >4 helices has yet to be achieved. Thus, we set out to design a heterohexameric, A_3_B_3_-type αHB for which the individual helices were unfolded in solution and co-assembled when mixed. As detailed below, our starting point was a *de novo* designed all-parallel, homohexameric αHB, CC-Hex2, for which we have sequence-to-structure relationships and an X-ray crystal structure.^8^

**Figure 1.**
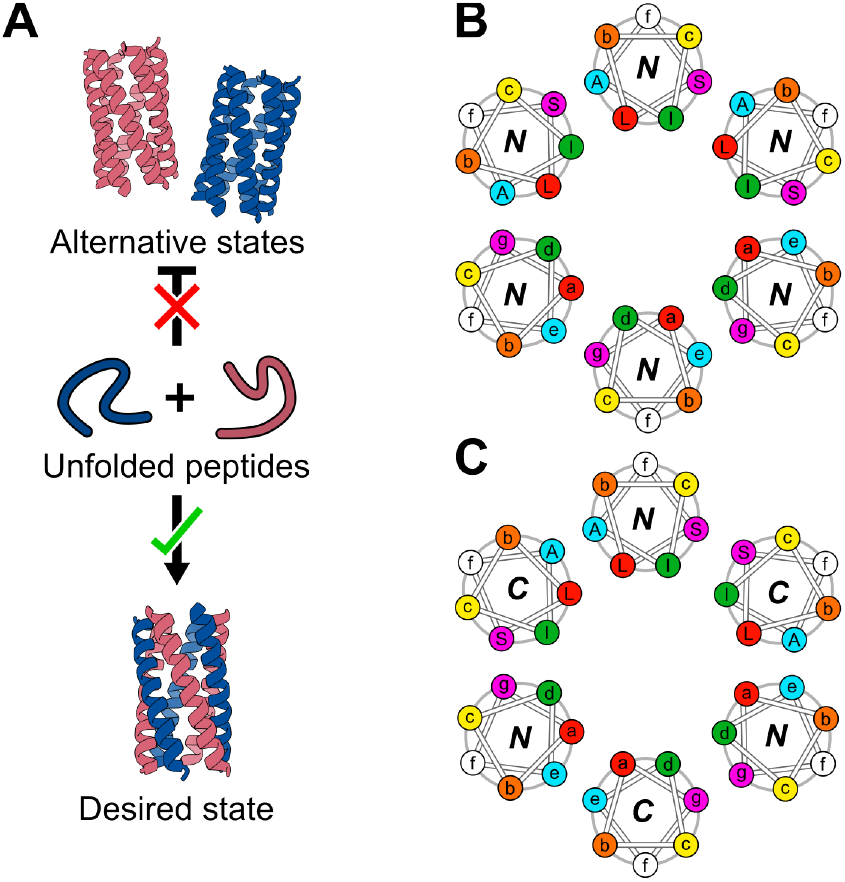
Design goal and features of coiled coils. (A) Our proposed design incorporates two differently charged peptides—an acidic peptide (red) and a basic peptide (blue)—that should not self-associate but fold only when both peptides are present into a 6-helix barrel. (B) Helical-wheel diagram for a 6-helix barrel with parallel helix orientations. Here, the ***a*** sites (and hence all heptad positions) are aligned with themselves on the *Z*-axis. (C) Helical-wheel diagram of a 6-helix barrel with antiparallel helix orientations. Here the ***a*** sites are more closely aligned vertically to the ***d*** sites.

## RESULTS AND DISCUSSION

The first aim of this study was to generate pairs of peptides that were unfolded in solution individually but assembled into a heterohexameric α-helical barrel (αHB) when mixed. Therefore, as a starting point, we took the CC-Hex2 peptide sequence, which forms a parallel homohexameric αHB in solution and the crystal state.^8^ This peptide has 4 heptad repeats starting at a ***c*** position of the heptad repeat, the ***g, a, d***, and ***e*** sites are occupied by serine (Ser, S), leucine (Leu, L), isoleucine (Ile, I) and alanine (Ala, A), respectively, which define the interhelical contacts and specify oligomeric state. Complementary charged glutamic acid (Glu, E) and lysine (Lys, K) residues at the flanking ***c*** and ***b*** sites, respectively, contribute to the helix-helix interactions through potential salt-bridge formation (Figure 1B).^8^ To design CC-Hex2-A_3_B_3_, we kept the ***g*-*a*-*d*-*e*** signature, and altered the peripheral ***b*** and ***c*** sites to generate acidic (A) and basic (B) peptides: in CC-Hex2-A, ***b*** = ***c*** = Glu; and in CC-Hex2-B, ***b*** = ***c*** = Lys.^4^ The remaining ***f*** positions were made to be glutamine (Gln, Q), Lys, and tryptophan (Trp, W) or tyrosine (Tyr, Y) for solubility and to add chromophores (Table 1). The 4-heptad peptides (Table S1) were made by solid-phase peptide synthesis (SPPS) and their identity confirmed by MALDI-TOF mass spectrometry (Figures S1 and S2).

**Table 1.**
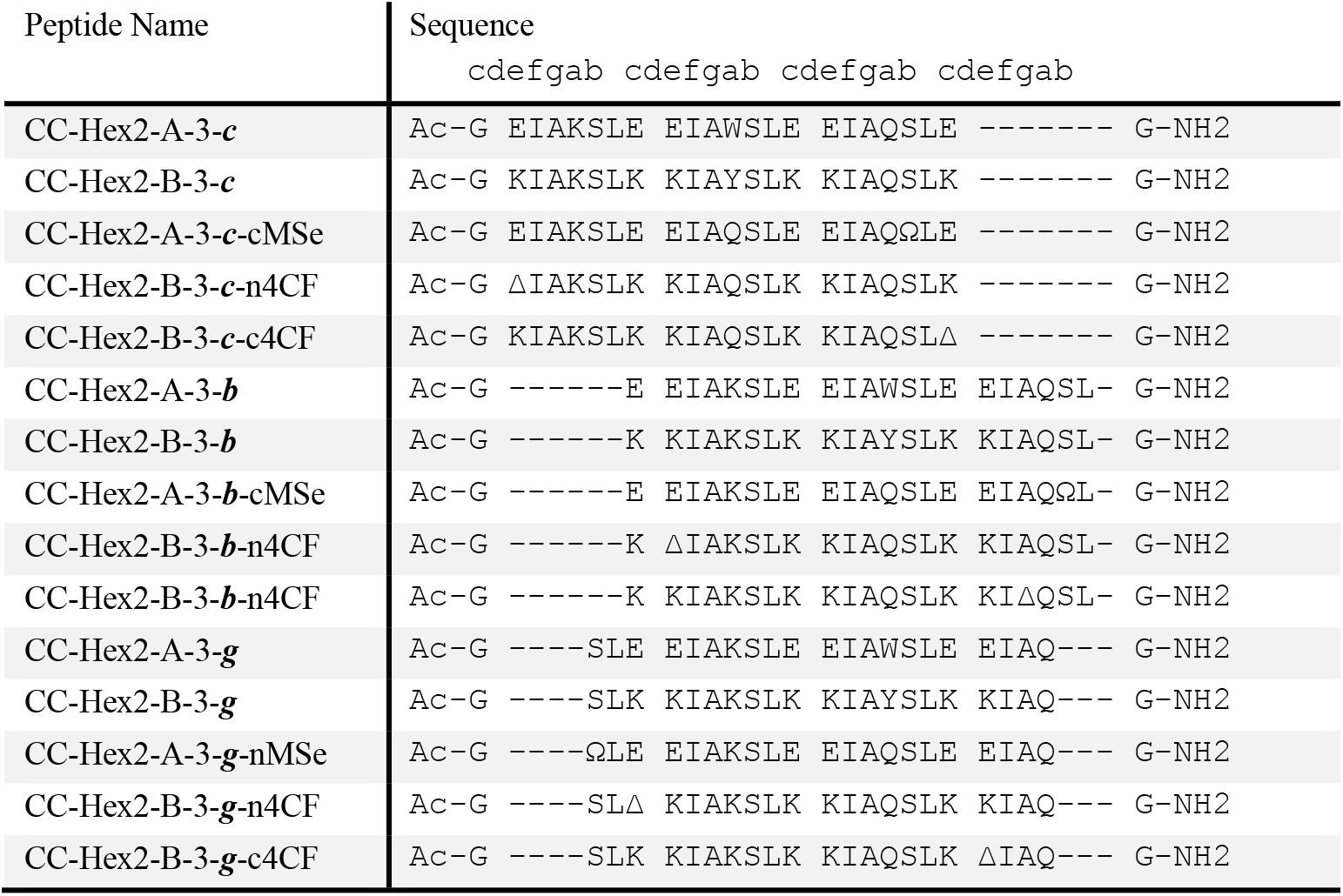
Designed sequences and summary of biophysical data for the peptides designed in this study. Δ represents 4-cyanophenylalanine (4CF) and Ω denotes selenomethionine (MSe). Ac-denotes acetylated *N*-terminal, -NH_2_ *C*-terminal amide groups.

To determine whether the peptides assembled conditionally, first we used circular dichroism (CD) spectroscopy to assess the secondary structure of the peptides in aqueous buffer at neutral pH (Figure S3). However, it quickly became apparent that our initial sequences did not meet the first of the design criteria, as the individual peptides formed highly α-helical structures and, moreover, precipitated when the anionic and cationic peptides were mixed. We have observed similar behavior with heterotetramers,^10^ and suggest that the highly and oppositely charged homomers form aggregates.

To reduce homo-oligomerization of CC-Hex2-A and CC-Hex2-B, we removed a heptad repeat from each sequence to reduce the helix-helix hydrophobic interactions and hence the stabilities of the target and off-target assemblies (Table 1). We called these peptides CC-Hex2-A-3-***c*** and CC-Hex2-B-3-***c***; A and B for the acidic and basic chains, 3 for the number of heptads, and ***c*** for the starting register (Table 1 and Figures S1 and S2). In this case, CD spectroscopy showed that the individual peptides were largely unfolded but formed a soluble and highly α-helical assembly when mixed (Figure 2A). The unfolded nature of the individual peptides and stability of the mixture were confirmed by variable-temperature CD measurements, which showed that the signals for CC-Hex2-A-3-***c*** and CC-Hex2-B-3-***c*** changed little upon heating, but that for the mixture revealed a thermally stable complex with the start of an unfolding transition at ≈ 80 ≈C, Figure 2B.

**Figure 2.**
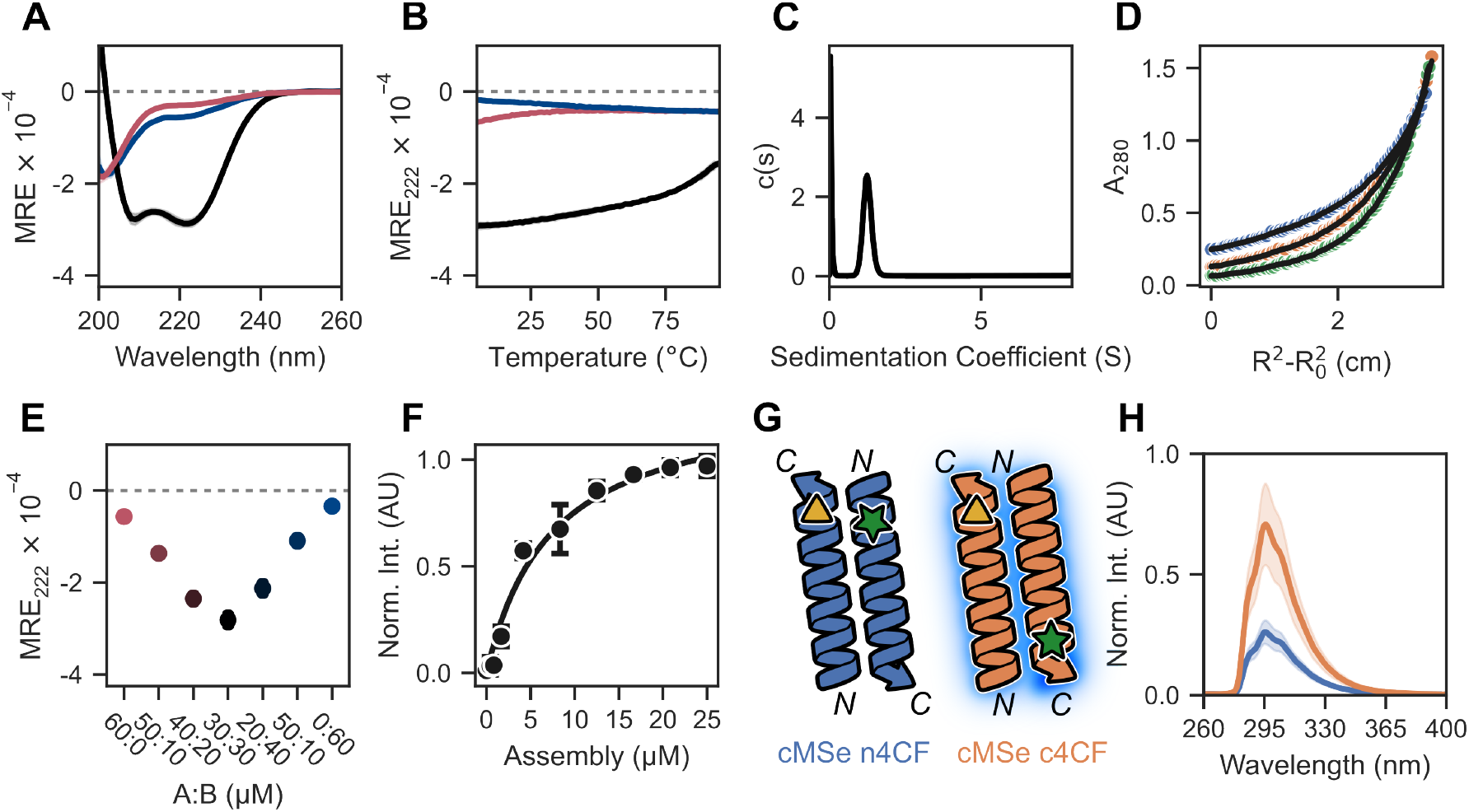
Biophysical characterization of CC-Hex2-AB-3-*c*.(A & B) CD spectra at 20 °C (A), and temperature-dependent CD signals monitored at 222 nm (B) for CC-Hex2-A-3-***c*** (red), CC-Hex2-B-3-***c*** (blue), and the mixture (black). (C) Analytical ultracentrifugation sedimentation velocity c(s) distribution fit recorded at 20 °C, 280 nm and 50 krpm 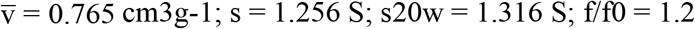.(D) Analytical ultracentrifugation sedimentation equilibrium data for CC-Hex2-AB-3-***c*** (fitted molecular weight = 15206 Da (6.0 x monomer mass), and 95% confidence limits = 14941 – 15502 Da. Speeds: 25 (blue), 35 (orange) 45 (green) krpm. (E) Concentration-dependent CD signal monitored at 222 nm for 60 μM CC-Hex2-AB-3-***c*** with different compositions of CC-Hex2-A-3-***c*** and CC-Hex2-B-3-***c***. Markers show the mean of the data and range bars represent one standard deviation; *N* = 3. (F) Saturation binding curves with DPH (1 μM); experimental data (filled circles) and fit (solid line) returns K_D_ = 7.3 ± 0.2 μM (R^2^ = 0.981). Peptide concentration was converted to αHB concentration assuming an oligomeric state of 6. (G) Cartoon depicting possible orientations of adjacent A-B helices in the proposed antiparallel complexes, and their predicted effects on the proximity of the 4-cyanophenylalanine fluorophore (4CF, green star) and the selenomethionine quencher (MSe, yellow triangle). ‘n’ and ‘c’ indicate mutations near the N and C termini, respectively. Fluorescence is quenched if the pair are adjacent. (H) Fluorescence quenching assay for labelled CC-Hex2-AB-3-***c*** peptides. CC-Hex2-A-3-***c***-cMSe + CC-Hex2-B-3-***c***-n4CF (blue) is quenched relative to CC-Hex2-A-3-***c***-cMSe + CC-Hex2-B-3-***c***-c4CF (orange). Filled regions represent one standard deviation of the mean; *N*=3.

Next, sedimentation velocity (SV) and sedimentation equilibrium (SE) analytical ultracentrifugation (AUC) experiments were used to probe the molecular weight of the A:B complex in solution. SV-AUC revealed a monodisperse oligomer of molecular weight ≈ 6 × the average monomeric masses of CC-Hex2-A-3-***c*** and CC-Hex2B-3-***c*** (Figure 2C), which was confirmed by SE-AUC (Figure 2D). We then determined the stoichiometry of the complex by monitoring the α-helical CD signal at 222 nm at different compositions of CC-Hex2-A-3-***c*** and CC-Hex2-B-3-***c*** (Figure 2E). This gave maximum signal at equimolar peptide concentrations, indicating a 1:1 stoichiometric complex. Combined, these data strongly indicated that we had made an A_3_B_3_ heterohexamer.

Coiled-coil assemblies above pentamer can form α-helical barrels with accessible central channels.^31^ To test whether we had made such a barrel—as opposed to a helical bundle with a consolidated hydrophobic core—the CC-Hex2-AB-3-***c*** assembly was tested with the environmentally sensitive dye, 1,3,5-diphenylhexatriene (DPH). When added to the peptide assembly, the dye fluoresced, and the concentration dependence of the response fitted to a single-site binding model returning a K_D_ of ≈ 7 μM (Figure 2F). This binding constant is similar to that for CC-Hex2^31^ and, thus, is indicative of CC-Hex2-AB-3-***c*** forming an open α-helical barrel.

Despite multiple attempts, we could not obtain diffraction-quality crystals for CC-Hex2-AB-3-***c*** mixtures needed for X-ray protein crystallography, and to determine directly whether the hexamer was parallel or antiparallel. Therefore, we used the proximity-based fluorescence-quenching assay illustrated in Figure 2G.^10, 32, 33^ The quencher selenomethionine (MSe, Ω) was incorporated at the *C*-terminal ***g*** site of CC-Hex2-A-3-***c***, and the fluorescent 4-cyanophenylalanine (4CF, Δ) was placed at either the *N*-terminal ***c*** site or the *C*-terminal ***b*** site of CC-Hex2-B-3-***c*** (Table 1). If the peptides assembled into a parallel structure, like the parent CC-Hex2, the combination with both *C-*terminal substitutions would lead to quenching of 4CF fluorescence, and, conversely, the *C-* plus *N-* combination would fluoresce. However, we found that the latter—*i*.*e*., the combination of the *C-*terminal quencher and the *N-*terminal fluorophore— quenched the fluorescence (Figure 2G, H). This indicated that the CC-Hex2-AB-3-***c*** complex was most likely an antiparallel assembly in solution.

This result was surprising as we had based the heterohexamer design on the CC-Hex2 sequence, which forms an all-parallel-helical assembly, confirmed by X-ray crystallography.^8^ However, it appears that, at least when reduced to 3 heptads in length and reconfigured into an A_3_B_3_ heteromeric design, specificity for parallel helix orientation is lost in favor of the antiparallel arrangement of helices (Figure 1B,C). Whilst parallel CCs are largely C_n_ symmetric structures in which the helices and their registers are fully aligned—*i*.*e*., the equivalent ***a*** sites of all helices are at the same level along the central *Z*-axis—many antiparallel structures have a helical offset (*Z*-shift) between adjacent helices (Figure 3A).^34^ This offset is believed to help with side-chain interdigitation to achieve better knobs-into-holes (KIH) packing of coiled coils.^5, 34-37^ In turn, this takes the sequences of adjacent helices out of alignment; *i*.*e*., ***a*** sites Cα atoms do not align with ***a*** site Cα atoms of adjacent helices, and so on.

**Figure 3.**
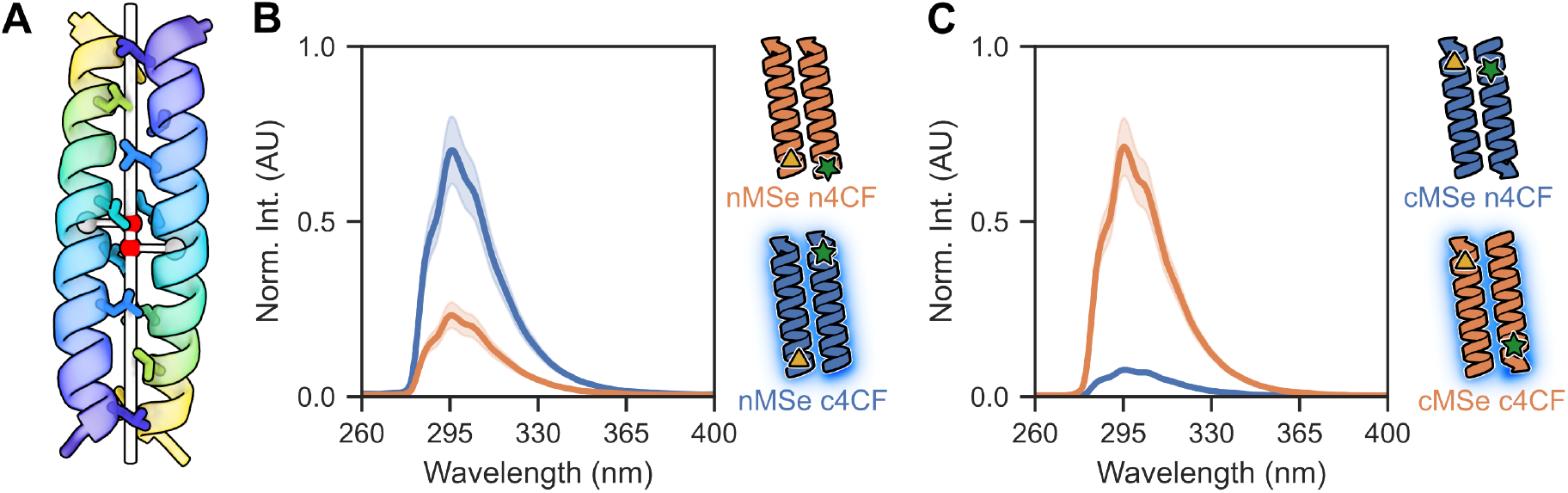
Manipulating helix orientation. (A) Antiparallel coiled coils slide along their *Z*-axes to facilitate interdigitation of side chains and KIH packing (sticks). The *Z*-shift is calculated as the distance between the projections of the centroids of each helix (gray spheres) onto the super helical axis (red spheres). For demonstration an antiparallel dimer is shown (PDB ID: 7Q1T). (B & C) Fluorescence quenching assays for labelled CC-Hex2-AB-3-***g*** (B) and CC-Hex2-AB-3-***b*** (C) peptides. The colored labels denote the positions of the 4CF and MSe pairs at the “n” or “c” termini.

On this basis, we wondered if the starting register of coiled-coil regions in peptide sequences—*i*.*e*., ***a, b, c, d, e, f*** or ***g***—might influence whether a *Z*-shift was accessible or not and hence the balance between parallel and antiparallel arrangements of helices. Therefore, to explore this, we synthesized peptides in the other six registers, i.e. CCHex2-AB-3-***a***, *-****b***, *-****d***, *-****e***, *-* ***f***, and *-****g*** and tested them experimentally (Table 1).

CD spectroscopy (Figure S3 and S4) and AUC experiments (Figure S9 and S10) were conducted for all new peptide variants (Table 1). Like the parent CCHex2-AB-3-***c***, CD spectroscopy showed that for CCHex2-AB-3-***a***, *-****b***, *-****d*** and -***g***, the individual A and B peptides were unfolded, but when mixed formed soluble and highly helical assemblies. In contrast, CCHex2-AB-3-***e*** and ***f*** aggregated and precipitated when the A and B peptides were mixed, suggesting the formation of aggregates rather than discrete assemblies. These were not pursued further in this study. As with CCHex2-AB-3-***c***, CCHex2-AB-3-***a***, *-****b, -d***, and *-****g*** were thermally stable, and showed the beginnings of unfolding transitions from ≈ 80 °C. In SV-AUC and SE-AUC experiments, CCHex2-AB-3-***a*** and -***d*** gave polydisperse assemblies ranging from 5 – 12 × the average monomeric masses. Therefore, these two systems were not explored further. AUC showed both CCHex2-AB-3-***b*** and -***g*** assemble into monodisperse hexamers. Concentrationdependent CD experiments confirmed A_3_:B_3_ stoichiometry (Figure S6 and S8), and both bound DPH with low µM affinities (Figure S11).

Next, we synthesized the chromophore- and quencher-labelled variants of the -***b*** and -***g*** pairs (Table 1, Figure 3) and mixed the different peptide combinations as described for CCHex2-AB-3-***c***. CC-Hex2-AB-3-***g*** showed reduced fluorescence when the substituted amino acids were positioned at the same terminus (*N-+ N-*), suggesting an assembly with a parallel helix orientation (Figure 3B). In contrast, CC-Hex2-AB-3-***b*** showed reduced fluorescence only when the residues were placed at different termini (*C-+ N-*, Figure 3C), suggesting an antiparallel assembly, like CC-Hex2-AB-3-***c***.

With a full set of solution-phase data for the two new peptide assemblies, we sought to determine X-ray structures for CC-Hex2-AB-3-***b*** and CC-Hex2-AB-3-***g***. Only CC-Hex2-AB-3-***g*** generated a usable diffraction pattern. The 1.9 Å resolution structure revealed a parallel heterohexameric coiled-coil barrel with alternating A and B chains (Figure 4A), which is completely consistent with the solution-phase data.

**Figure 4.**
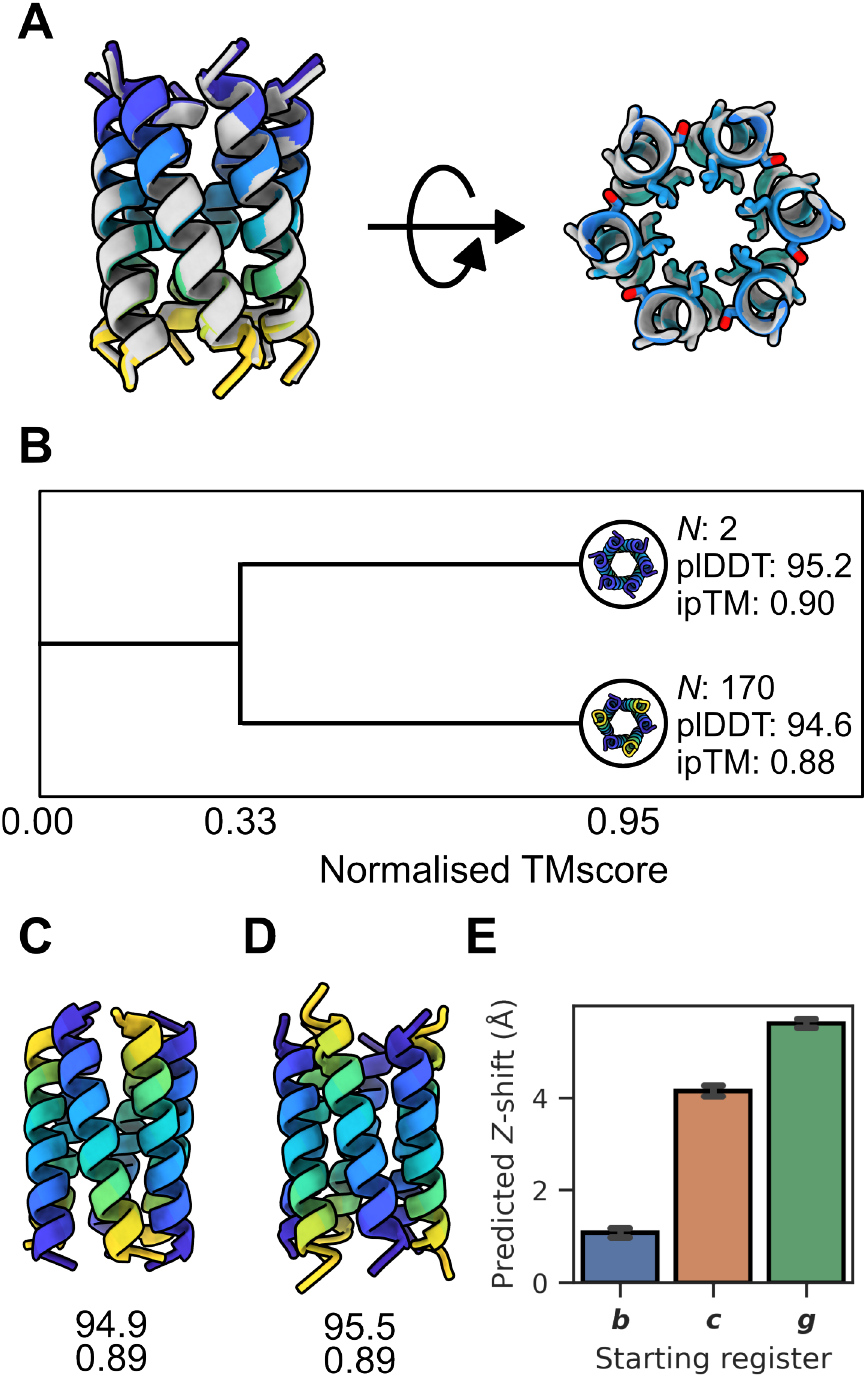
Structural features of CC-Hex2-AB-3-*b*/*c*/*g*.(A) 1.9 Å X-ray crystal structure of CC-Hex2-AB-3-***g*** (PDB ID: 9EVG, grey) aligned to its top ranked computed structure model (colored, Cα-RMSD: 0.52 Å). Left, side on view of the all-parallel arrangement shows a *Z*-shift near 0 Å between helices. Right, knob-into-hole packing. For clarity, only the Cα-Cβ bonds are shown for side chains at the ***b, c***, and ***f*** positions. (B) Clustering of the best AF-M predictions (ipTM > 0.7). Clustered by normalized TMscore (structure similarity = 0.95). Two clusters appear, representing the parallel and antiparallel predictions. Representatives are taken as the model with the highest confidence metrics. *N* denotes cluster size. Bottom, top ranked computed structure model of CC-Hex2-AB-3-***b*** (C) and CC-Hex2-AB-3-***c*** (D). Numbers denote plDDT (top) and ipTM (bottom). (E) AF-M computed structure model predicted *Z*-shift between helices of antiparallel poses.

In lieu of experimental atomic-resolution structures for CC-Hex2-AB-3-***b*** and ***c***, we computed structural models with AlphaFold-MultimerV3 (AF-M) implemented in ColabFold.^38, 39^ First, we tested its capability to predict the CC-Hex2-AB-3-***g*** structure. Initially, the default configuration of ColabFold was used, which only sampled a small degree of model diversity with 1 seed used for each of the 5 AF-M models, creating a total of 5 predictions. With this limited sampling, the top prediction was highly confident (ipTM > 0.7), but it did not match the crystal structure: the experimental structure is all parallel, but the model predicted antiparallel helices (Figure 4A). Others have shown that more extensive sampling may be required to correctly predict multimeric structures.^38, 40^ Therefore, we increased sampling through additional random seeds. This alters the stochastic elements of the model, which can lead to more model diversity and consequently better predictions. We increased the number of random seeds from 1 to 100, generating 100 predictions per model for a total of 500 predictions per sequence. To observe the structural diversity of the predictions, high confidence results (ipTM > 0.7, *N* = 172) were clustered by structural similarity using a normalized TMscore (TMscore × fraction aligned, Figure 4B).^41^ This gave two groups when clustered by a structure similarity of 0.95 with all-parallel (*N* = 2) and antiparallel (*N* = 170) arrangements of helices. Thus, overall, the AlphaFold predictions were biased towards the incorrect antiparallel models by 85:1, which is concerning. That said, the 2 highest-confidence predictions were for the parallel models (Figure 4B). Indeed, these were near-superposable with the X-ray crystal structure of CC-Hex2-AB-3-***g*** (Cα-RMSD = 0.52 Å, Figure 4A).

Mindful of the caveat in this approach, we applied it to generate models for CC-Hex2-AB-3-***b*** and ***c***, again using 100 seeds across the 5 different modes for 500 total predictions each. The ratios of predicted models with all-parallel and with antiparallel arrangements of helices were 1:301 and 1:115, respectively. However, the top-ranked predictions for both sequences were antiparallel 6-helix barrels with alternating A and B chains consistent with the solution-phase data (Figure 4C, D). On this basis, we assumed that these top predicted models for CCHex2-AB-3-***b*** and CCHex2-AB-3-***c*** might best represent the actual structures. With these models and the “theoretical”, but not experimentally observed, model of the antiparallel state of CCHex2-AB-3-***g***, we examined the hypothesis of *Z*-shift driving helix orientation. Z-shifts of the high-confidence models (ipTM > 0.7) for the three registers were calculated (Figure 4E). Interestingly, these were in the order ***g*** > ***c*** > ***b***. Moreover, at 5.6 Å, the *Z*-shift for the ***-g*** peptide is a whole α-helical turn (5.4 Å). Reasoning that larger *Z*-shifts lead to larger helical overhangs and less helix-helix contacts in the assembled CCs, it makes sense that the antiparallel arrangement for the ***g-***register peptide would be the least stable of these, and less stable than the observed parallel structure. This analysis suggests a *Z*-shift threshold for the CC-Hex2-AB-3 designs to form antiparallel assemblies of ≈ 4 Å, and above this the parallel state is favored. Although our analysis is limited and focused, it fits with larger analyses of experimental structures of parallel and antiparallel CCs from the RCSB PDB, albeit for smaller oligomers.^34, 42^ From these studies, parallel CCs have *Z*-shift values (referred to as Δ*Z* and axial shift, and defined slightly differently in those papers) are sharply centered around 0; whereas in antiparallel structures, they are ≈ 2 – 3 Å.

## CONCLUSION

Through a combination of rational peptide design and empirical redesign, we have delivered three heterohexameric A_3_B_3_-type α-helical barrel assemblies (αHBs). One of these, CC-Hex2-AB-3-***g***, has all-parallel helices as is confirmed by solution-phase spectroscopic data and an X-ray peptide crystal structure. The data for the other two, CC-Hex2-AB-3-***b*** and CC-Hex2-AB-3-***c***, are consistent with antiparallel arrangements of helices in solution, but we could not confirm this by X-ray crystallography. We find that the sequences can be modelled as the parallel and antiparallel states by AlphaFold-Multimer (AF-M). However, this requires larger sampling than offered by the default AlphaFold settings, and then close inspection of the models and prediction metrics. This is because, at least for our system, AF-M appears biased to predict antiparallel helix orientations. This is opposite to other observations for lower oligomeric-state coiled coils (< 5), which report a bias for parallel predictions.^43^

The serendipitous discovery that CC-Hex2-AB-3-***c*** forms an antiparallel assembly led to the hypothesis that the starting register of the coiled-coil sequence in peptide designs could contribute to specifying coiled-coil topology, (*i*.*e*., parallel, antiparallel, or mixed arrangements of helices. The reasoning is that different starting registers would lead to different helical overhangs for essentially the same knobs-into-hole packing between helices; and the longer the overhang the less stable the antiparallel state relative to the parallel state, where the helices are usually aligned and do not overhang. To explore this, we examine the *Z*-shift of AF-M models for the antiparallel states of the ***b-, c-*** and ***g-***register peptides for which we have experimental data. From this, the antiparallel model for CC-Hex-2-AB-***g*** has the largest predicted *Z*-shift and this is outside the range observed across structurally defined natural CCs, albeit of lower oligomeric states.^34, 42^ This is consistent with that peptide forming a parallel rather than an antiparallel assembly. This adds another design parameter that should be considered for controlling helix orientation in *de novo* coiled-coil peptides assemblies and proteins.

This work adds to existing literature on understanding coiled-coil peptide design and assembly.^10, 29, 30, 44^ To date, the coiled coil-design hierarchy has been: First, focus on residues at the largely hydrophobic, knobs-into-holes, helix-helix interfaces, as these have most influence on coiled-coil stability and oligomeric state. Second, use inter-helical salt-bridges to direct coiled-coil partnering, *i*.*e*. to form homo-or heteromeric assemblies. However, these can be used to steer parallel versus antiparallel helix orientations with what might be termed “classical” and “bar-magnet” charge patterning, respectively.^10, 11^ Here we add sequence register to the list of features that can influence coiled-coil assembly and helix orientation. However, we recognize that its contribution may be small. For example, here we show that CC-Hex2-AB-3-***c*** forms antiparallel A_3_B_3_-type assembly in solution. Nonetheless, most previously reported *de novo* αHBs have ***c***-register and are parallel assemblies.^8, 9^ Similarly, while CC-Hex2-AB-3-***g*** forms a parallel A_3_B_3_ αHB, the antiparallel tetramer, ap-CC-Tet, also has a ***g***-register but with bar-magnet charge patterning to drive the antiparallel state.^10^ Nonetheless, we suggest that sequence register should be considered as a variable when assessing new coiled coil-based peptide and protein designs, as its subtle effects may be important particularly as peptide and protein design move on from targeting static structures to more-dynamic and functional assemblies.

## ASSOCIATED CONTENT

**Supporting Information**

The following file is available free of charge.

Materials and methods; supplementary data for all peptides investigated in study (file type, PDF)

Accession Codes CC-Hex2-AB-3-***c***, PDB ID: 9EVG

## AUTHOR INFORMATION

### Author Contributions

J.J.C, K.I.A, A.R, and D.N.W conceived the study and contributed to the experimental design. J.J.C and K.I.A designed and characterized the peptides. K.I.A solved the X-ray crystal structure. J.J.C conducted the computational studies. J.J.C, K.I.A, A.R and D.N.W wrote the paper.

### Notes

The authors declare no competing financial interest.

## ACKNOWLEDGMENT

J.J.C and D.N.W were supported by the Engineering and Physical Sciences Research Council (EP/R513179/1). J.J.C and A.R were also supported by the Australian Research Council Industrial Transformation Training Centre for Facilitated Advancement of Australia’s Bioactives (Grant IC210100040). We are also grateful to the Max Planck-Bristol Centre for Minimal Biology, which supports K.I.A. and D.N.W. We thank the Mass Spectrometry Facility, School of Chemistry, University of Bristol, for access to the EPSRC-funded Bruker Ultraflex MALDI–TOF instrument (EP/K03927X/1). We would like to thank Diamond Light Source for access to the I24 beamlines (proposals mx2373). Finally, we thank A. Boyle for helping proofread the manuscript.

## Notes

### Competing Interest Statement

The authors have declared no competing interest.

## REFERENCES

(1) Korendovych, I. V., DeGrado, W. F. (2020) De novo protein design, a retrospective. Q. Rev. Biophys. 53, e3.

(2) Woolfson, D. N. (2021) A Brief History of De Novo Protein Design: Minimal, Rational, and Computational. J. Mol. Biol. 433, 167160.

(3) Kortemme, T. (2024) De novo protein design-From new structures to programmable functions. Cell 187, 526–544.

(4) Woolfson, D. N. (2023) Understanding a protein fold: The physics, chemistry, and biology of α-helical coiled coils. J. Biol. Chem. 299, 104579.

(5) Lupas, A. N., Bassler, J. (2017) Coiled Coils - A Model System for the 21st Century. Trends Biochem. Sci. 42, 130–140.

(6) Squire, J. M., Parry, D. A. (2017) Fibrous Protein Structures: Hierarchy, History and Heroes. Subcell. Biochem. 82, 1–33.

(7) Fletcher, J. M., Boyle, A. L., Bruning, M., Bartlett, G. J., Vincent, T. L., Zaccai, N. R., Armstrong, C. T., Bromley, E. H., Booth, P. J., Brady, R. L., et al. (2012) A basis set of de novo coiled-coil peptide oligomers for rational protein design and synthetic biology. ACS Synth. Biol. 1, 240–250.

(8) Thomson, A. R., Wood, C. W., Burton, A. J., Bartlett, G. J., Sessions, R. B., Brady, R. L., Woolfson, D. N. (2014) Computational design of water-soluble α-helical barrels. Science 346, 485–488.

(9) Dawson, W. M., Martin, F. J. O., Rhys, G. G., Shelley, K. L., Brady, R. L., Woolfson, D. N. (2021) Coiled coils 9-to-5: rational de novo design of α-helical barrels with tunable oligomeric states. Chem. Sci. 12, 6923–6928.

(10) Naudin, E. A., Albanese, K. I., Smith, A. J., Mylemans, B., Baker, E. G., Weiner, O. D., Andrews, D. M., Tigue, N., Savery, N. J., Woolfson, D. N. (2022) From peptides to proteins: coiled-coil tetramers to single-chain 4-helix bundles. Chem. Sci. 13, 11330–11340.

(11) Albanese, K. I., Petrenas, R., Pirro, F., Naudin, E. A., Borucu, U., Dawson, W. M., Scott, D. A., Leggett, G. J., Weiner, O. D., Oliver, T. A. A., et al. (2024) Rationally seeded computational protein design of ɑ-helical barrels. Nat. Chem. Biol. 20, 991–999.

(12) Crick, F. H. C. (1953) The packing of α-helices: simple coiled-coils. Acta Crystallogr. 6, 689–697.

(13) Crick, F. H. C. (1953) The Fourier transform of a coiled-coil. Acta Crystallogr. 6, 685–689.

(14) Pauling, L., Corey, R. B. (1953) Compound helical configurations of polypeptide chains: structure of proteins of the alpha-keratin type. Nature 171, 59–61.

(15) Huang, P. S., Oberdorfer, G., Xu, C., Pei, X. Y., Nannenga, B. L., Rogers, J. M., DiMaio, F., Gonen, T., Luisi, B., Baker, D. (2014) High thermodynamic stability of parametrically designed helical bundles. Science 346, 481–485.

(16) Burton, A. J., Thomson, A. R., Dawson, W. M., Brady, R. L., Woolfson, D. N. (2016) Installing hydrolytic activity into a completely de novo protein framework. Nat. Chem. 8, 837–844.

(17) Dawson, W. M., Shelley, K. L., Fletcher, J. M., Scott, D. A., Lombardi, L., Rhys, G. G., LaGambina, T. J., Obst, U., Burton, A. J., Cross, J. A., et al. (2023) Differential sensing with arrays of de novo designed peptide assemblies. Nat. Commun. 14, 383.

(18) Scott, A. J., Niitsu, A., Kratochvil, H. T., Lang, E. J. M., Sengel, J. T., Dawson, W. M., Mahendran, K. R., Mravic, M., Thomson, A. R., Brady, R. L., et al. (2021) Constructing ion channels from water-soluble α-helical barrels. Nat. Chem. 13, 643–650.

(19) O’Shea, E. K., Lumb, K. J., Kim, P. S. (1993) Peptide ‘Velcro’: design of a heterodimeric coiled coil. Curr. Biol. 3, 658–667.

(20) Nautiyal, S., Woolfson, D. N., King, D. S., Alber, T. (1995) A designed heterotrimeric coiled coil. Biochemistry 34, 11645–11651.

(21) Fairman, R., Chao, H. G., Lavoie, T. B., Villafranca, J. J., Matsueda, G. R., Novotny, J. (1996) Design of heterotetrameric coiled coils: evidence for increased stabilization by Glu(-)-Lys(+) ion pair interactions. Biochemistry 35, 2824–2829.

(22) Nautiyal, S., Alber, T. (1999) Crystal structure of a designed, thermostable, heterotrimeric coiled coil. Prot. Sci. 8, 84–90.

(23) Thomas, F., Boyle, A. L., Burton, A. J., Woolfson, D. N. (2013) A Set of de Novo Designed Parallel Heterodimeric Coiled Coils with Quantified Dissociation Constants in the Micromolar to Sub-nanomolar Regime. J. Am. Chem. Soc. 135, 5161–5166.

(24) Bermeo, S., Favor, A., Chang, Y. T., Norris, A., Boyken, S. E., Hsia, Y., Haddox, H. K., Xu, C., Brunette, T. J., Wysocki, V. H., et al. (2022) De novo design of obligate ABC-type heterotrimeric proteins. Nat. Struct. Mol. Biol. 29, 1266–1276.

(25) Kashiwada, A., Hiroaki, H., Kohda, D., Nango, M., Tanaka, T. (1999) Design of a Heterotrimeric α-Helical Bundle by Hydrophobic Core Engineering. J. Am. Chem. Soc. 122, 212–215.

(26) Lebruin, L. T., Banerjee, S., O’Rourke, B. D., Case, M. A. (2011) Metal ion-assembled micro-collagen heterotrimers. Biopolymers 95, 792–800.

(27) Tolbert, A. E., Ervin, C. S., Ruckthong, L., Paul, T. J., Jayasinghe-Arachchige, V. M., Neupane, K. P., Stuckey, J. A., Prabhakar, R., Pecoraro, V. L. (2020) Heteromeric three-stranded coiled coils designed using a Pb(II)(Cys)3 template mediated strategy. Nat. Chem. 12, 405–411.

(28) Zaccai, N. R., Chi, B., Thomson, A. R., Boyle, A. L., Bartlett, G. J., Bruning, M., Linden, N., Sessions, R. B., Booth, P. J., Brady, R. L., et al. (2011) A de novo peptide hexamer with a mutable channel. Nat. Chem. Biol. 7, 935–941.

(29) Spencer, R. K., Hochbaum, A. I. (2016) X-ray Crystallographic Structure and Solution Behavior of an Antiparallel Coiled-Coil Hexamer Formed by de Novo Peptides. Biochemistry 55, 3214–3223.

(30) Spencer, R. K., Hochbaum, A. I. (2017) The Phe-Ile Zipper: A Specific Interaction Motif Drives Antiparallel Coiled-Coil Hexamer Formation. Biochemistry 56, 5300–5308.

(31) Thomas, F., Dawson, W. M., Lang, E. J. M., Burton, A. J., Bartlett, G. J., Rhys, G. G., Mulholland, A. J., Woolfson, D. N. (2018) De Novo-Designed α-Helical Barrels as Receptors for Small Molecules. ACS Synth. Biol. 7, 1808–1816.

(32) Rhys, G. G., Cross, J. A., Dawson, W. M., Thompson, H. F., Shanmugaratnam, S., Savery, N. J., Dodding, M. P., Höcker, B., Woolfson, D. N. (2022) De novo designed peptides for cellular delivery and subcellular localisation. Nat. Chem. Biol. 18, 999–1004.

(33) Watson, M. D., Peran, I., Raleigh, D. P. (2016) A Non-perturbing Probe of Coiled Coil Formation Based on Electron Transfer Mediated Fluorescence Quenching. Biochemistry 55, 3685–3691.

(34) Grigoryan, G., DeGrado, W. F. (2011) Probing Designability via a Generalized Model of Helical Bundle Geometry. J. Mol. Biol. 405, 1079–1100.

(35) Rhys, G. G., Wood, C. W., Beesley, J. L., Zaccai, N. R., Burton, A. J., Brady, R. L., Thomson, A. R., Woolfson, D. N. (2019) Navigating the Structural Landscape of De Novo α-Helical Bundles. J. Am. Chem. Soc. 141, 8787–8797.

(36) Gernert, K. M., Surles, M. C., Labean, T. H., Richardson, J. S., Richardson, D. C. (1995) The Alacoil: a very tight, antiparallel coiled-coil of helices. Prot. Sci. 4, 2252–2260.

(37) Hill, R. B., Raleigh, D. P., Lombardi, A., DeGrado, W. F. (2000) De novo design of helical bundles as models for understanding protein folding and function. Acc. Chem. Res. 33, 745–754.

(38) Evans, R., O’Neill, M., Pritzel, A., Antropova, N., Senior, A., Green, T., Žídek, A., Bates, R., Blackwell, S., Yim, J., et al. (2022) Protein complex prediction with AlphaFold-Multimer. bioRxiv 2021.2010.2004.463034.

(39) Mirdita, M., Schütze, K., Moriwaki, Y., Heo, L., Ovchinnikov, S., Steinegger, M. (2022) ColabFold: making protein folding accessible to all. Nat. Methods 679–682.

(40) Wallner, B. (2023) AFsample: improving multimer prediction with AlphaFold using massive sampling. Bioinformatics 39.

(41) Zhang, Y., Skolnick, J. (2004) Scoring function for automated assessment of protein structure template quality. Proteins 57, 702–710.

(42) Dunin-Horkawicz, S., Lupas, A. N. (2010) Measuring the conformational space of square four-helical bundles with the program samCC. J. Struc. Biol. 170, 226–235.

(43) Madaj, R., Martinez-Goikoetxea, M., Kaminski, K., Ludwiczak, J., Dunin-Horkawicz, S. (2024) Applicability of AlphaFold2 in the modelling of coiled-coil domains. bioRxiv 2024.2003.2007.583852.

(44) Rhys, G. G., Dawson, W. M., Beesley, J. L., Martin, F. J. O., Brady, R. L., Thomson, A. R., Woolfson, D. N. (2021) How Coiled-Coil Assemblies Accommodate Multiple Aromatic Residues. Biomacromolecules 22, 2010–2019.

