## Supplementary information for "*De novo* design of parallel and antiparallel A_3_B_3_ heterohexameric α-helical barrels"

### 1 MATERIALS AND METHODS

#### 1.1 General

All reagents were purchased from Sigma Aldrich (Gillingham, UK), Fisher Scientific (Loughborough, UK) or Merck (Darmstadt, Germany) and used without further purification. All Fmoc amino acids were purchased from either Sigma Aldrich or Fluorochem. Biophysical data collection was typically carried out in HEPES buffered saline (HBS, 25 mM HEPES, 100 mM NaCl, pH 7.5, water). When variations of these are used, they are explicitly stated. Isoelectric points, masses, and molar extinction coefficients at 280 nm ($\varepsilon_{280\ nm}$) for peptides were calculated using Innovagen's peptide property calculator. Molar extinction coefficients for selected peptides at 214 nm ($\varepsilon_{214\ nm}$) were calculated individually based on their polypeptide sequence. Diagram representations protein/peptide structures and computed structure models were all processed in ChimeraX molecular visualization software. All instances of water refer to ultrapure Milli-Q™. All 96-well plates were dispensed using an Eppendorf epMotion® 5070 liquid handling robot (Hamburg, Germany). Pure peptides stocks were stored at -20 °C. All heteromeric peptide combinations were mixed in a 1:1 ratio unless otherwise specified.

#### 1.2 Automated Fmoc peptide synthesis

Automated microwave Solid-phase peptide synthesis (SPPS) was performed on a CEM Liberty Blue apparatus (Buckingham, UK) synthesizer with inline UV monitoring. Syntheses were performed on 0.1 mmol scales. The resin (Rink amide MBHA, 0.65 mmol/g loading, 100 – 200 mesh) was weighed to enable a synthesis on a 0.1 mmol scale. The peptide coupling reactions were performed by adding Fmoc protected amino acids dissolved in dimethylformamide (DMF) (2.5 mL, 0.2 M), the coupling reagent *N,N*-diisopropylcarbodiimide (DIC) in DMF (1.0 mL, 1 M) and Oxyma Pure in DMF (1.0 mL, 0.5 M) to the respective resin. Standard couplings were performed at 90 °C for 4.5 mins (100 W for 20 s, 60 W for 10 s, 35 W for 240 s). Standard deprotections were performed using 20% (v/v) morpholine in DMF at 90 °C for 1.5 mins (125 W 30 s, 32 W 60 s). All peptides were manually acetyl capped through addition of pyridine (0.5 mL) and acetic anhydride (0.25 mL) in DMF (9.25 mL), shaking at room temperature (rt) for 20 minutes. The resin was washed three time with DMF followed by three times with dichloromethane (DCM) before cleavage. Peptides were cleaved from the resin with addition of 5 mL of a mixture 95:2.5:2.5 v/v trifluoroacetic acid (TFA)/$H_2O$/triisopropylsilane (TIPS), shaking at room temperature for 3 hours. The TFA solution was then filtered to remove the resin beads and was reduced in volume to ≈5 mL or lower using a flow of $N_2$. Cleaved peptide was precipitated with cold diethyl ether (≈40 mL), isolated via centrifugation, and dissolved in a 1:1 mixture acetonitrile (MeCN)/$H_2O$. Crude peptides were lyophilized to yield a white or off-white powder.

#### 1.3 Semi-preparative High Performance Liquid Chromatography (HPLC)

All peptides were purified by reverse phase HPLC on JASCO apparatus fitted with pumps (PU-980), degasser (DG-980-50), UV/Vis detector (UV-2077) and column oven (CO-1560), controlled by a LC-NetII/ADC. HPLC was used with a Phenomenex Luna C18 (Macclesfield, UK) column (150 x 10 mm, 5 mm particle size, 100 Å pore size). Crude peptide was dissolved at 5 mg/mL in 20 or 40 % v/v MeCN in $H_2O$ with 0.1% TFA, injected to the column and eluted with a 3 mL/min linear gradient (20 or 40 – 100%) of MeCN in $H_2O$ with 0.1% TFA (pI > 7) or 25 mM ammonium bicarbonate (pI < 7) each over 30 minutes. Elution of the peptide was detected with inline UV monitoring at 220 and 280 nm wavelengths simultaneously. A column oven (50 °C) was employed to improve separation. Pure fractions were identified by analytical HPLC and matrix-assisted laser desorption/ionization–time of flight (MALDI-TOF) mass spectrometry, then pooled, and freeze-dried.

#### 1.4 Analytical HPLC

Analytical HPLC traces were obtained using a JASCO 2000 series HPLC system (Hachioji, Tokyo) and a Phenomenex Kinetex C18 (100 x 4.6 mm, 5 µm particle size, 100 Å pore size) column (Macclesfield, UK). Chromatograms were monitored at 220 and 280 nm wavelengths. The linear gradient matched that used for semi-preparative HPLC but over 25 min at a flow rate of 1 mL/min. When required, a JASCO column oven (CO-1650) was used at 50 °C was employed to assist peptide elution.

#### 1.5 Mass spectrometry

Matrix-assisted laser desorption/ionization–time of flight (MALDI-TOF) mass spectra were collected on a Bruker UltrafleXtreme MALDI-TOF mass spectrometer (Coventry, UK) operating in positive-ion reflector mode. Peptides were spotted on a ground steel target plate using α-cyano-4-hydroxycinnamic acid dissolved in 1:1 MeCN/$H_2O$ as the matrix. Masses quoted are for the monoisotopic mass as the singly protonated species.

#### 1.6 Peptide concentration determination

Peptide concentration was determined at 280 nm using a Thermo Scientific Nanodrop 2000 (Waltham, USA) spectrometer or an Agilent (Stockport, UK) Cary 100 UV/Vis spectrophotometer ($\varepsilon_{280}$(Trp) = 5690 $M^{-1}$ $cm^{-1}$; $\varepsilon_{280}$ (Tyr) = 1280 $M^{-1}$ $cm^{-1}$). For sequences without a Trp or Tyr, concentrations were determined using the amide absorbance for each sequence, using values determined by Gruppen and Kuipers using the equation[1]:

$$\varepsilon_{peptide}(M^{-1}cm^{-1}) = \varepsilon_{peptide\ bond} \times n_{peptide\ bonds} + \sum_{i=1}^{20} \varepsilon_{amino\ acid(i)} \times n_{amino\ acid(i)}$$

where $\varepsilon_{peptide\ bond}$ is 923 $M^{-1}$ $cm^{-1}$ and $\varepsilon_{amino\ acid}$ corresponds to the molar extinction coefficient of the free amino acid at 214 nm.

#### 1.7 Circular dichroism (CD) spectroscopy

Circular dichroism (CD) spectra were collected on JASCO (Hachioji, Tokyo) J-810 or J-815 spectrophotometers fitted with a Peltier temperature controller. Measurements were carried out in quartz cuvettes (Starna Scientific; Ilford, UK) with pathlength used dependent on peptide concentration (1 mm: > 50 µM; 5 mm: 10–50 µM; 10 mm: < 10 µM). The instrument was set with a sensitivity of 100 mdeg, a pitch and bandwidth of 1 nm and a scan rate of 100 nm.$min^{-1}$ at 20 °C with a response time of 1 s. Readings were averaged over 8 accumulative scans. Readings were taken from 200–260 nm. Peptide samples were made up in HBS. Baseline recordings using the same buffer, cuvette and instrument parameters were subtracted from each spectrum.

Temperature variable scans were taken at 222 nm ranging from 5 – 95 °C at a data pitch of 1 °C with a 16 s delay and a temperature ramping rate dependent on path length (1 mm: 60 °C.hr$^{-1}$, 5 mm: 45 °C.hr$^{-1}$, 10 mm: 30 °C.h$^{-1}$).

To standardize the data, each spectrum was converted from ellipticities (mdeg) to mean residue ellipticities (MRE, deg cm$^2$.dmol$^{-1}$.res$^{-1}$) by normalizing for the concentration of peptide bonds and the cuvette path lengths:

$$[\theta]\ (deg.cm^2.dmol^{-1}.res^{-1}) = \theta(mdeg)/c \times l \times n \times 10$$

where the variable $\theta$ is the measured difference in absorbed circularly polarized light in millidegrees, $c$ is the μM concentration of the compound, $l$ is the path length of the cuvette in mm, and $n$ is the number of amide bonds in the polypeptide, including the *N*-terminal acetyl.

**Analytical ultracentrifugation**

Analytical ultracentrifugation sedimentation velocity (SV) was completed at 20 °C in either a Beckman Optima XL-A or an XL-I analytical ultracentrifuges equipped with an An-50 Ti rotor. SV cells were constructed with either aluminum or epon 2-channel centerpieces and quartz windows (Beckman Coulter; High Wycombe, UK). Sample channels were made up of 300 or 400 μL of sample for epon and aluminum cells. Samples consisted of 150 μM peptide in HBS. The reference channels (epon, aluminum) contained 310 or 410 μL buffer. Samples were centrifuged at 50,000 rpm with absorbance scans (280 nm) taken across a radial range of 5.8–7.3 cm at 5 min intervals after an initial 5 min delay for 120 scans. The data acquired were fitted to a continuous c(s) distribution model using SEDFIT[2] with 95 % confidence limits. The baseline, bottom, frictional coefficient ($f/f_0$) and systematic time-invariant noise were fitted. SEDNTERP[2] was used to calculate partial specific volumes ($\bar{v}$) of peptides and buffer densities (ρ) and viscosities (η).

Analytical ultracentrifugation sedimentation equilibrium (SE) was conducted at 20 °C in a Beckman Optima XL-A or XL-I analytical ultracentrifuge equipped with an An-50 Ti rotor. The SE cells were composed of a 6-channel epon centerpiece with quartz windows. The sample channel was made up of 70 μM of peptide (35 μM each A and B peptides) in HBS.

The reference channels had 120 µL buffer. Absorbance scans (280 nm) were taken every 8 hours, with a second scan to monitor the equilibrium state taken 1 hour later at speeds between 20,000-48,000 rpm. Data were fitted to a single ideal species model using SEDPHAT[2]. Statistical analysis was carried out by a Monte Carlo analysis (1000 iterations, randomized start points, 95 % confidence limits) of acquired fits.

#### 1.8 DPH binding assay

All ligand binding fluorescence experiments were recorded in a BMG Labtech (Aylesbury, UK) Clariostar plate reader at 25 °C. Binding experiments were completed with 1,6-diphenyl-1,3,5-hexatriene (DPH) at 1 µM in HBS and 5 % v/v dimethyl sulfoxide (DMSO). Peptides were titrated into the DPH solutions at concentrations ranging from 1 – 300 µM. These were then calculated into relative assembly concentration by dividing by the oligomeric state. Peptide and ligand were equilibrated for 2 hours at rt with constant shaking. The fluorescence spectra were measured using excitation ($\lambda_{ex}$) of 1,6-diphenyl-1,3,5-hexatriene (DPH) with emissions ($\lambda_{em}$) at 455 nm (± 10 nm). Dissociation constants ($K_d$) were calculated by fitting a single-site binding model:

$$y = \frac{B_{max} \cdot x}{K_D + x}$$

where $x$ is the concentration of peptide, $B_{max}$ is the fluorescence signal when all the constant component is bound, and $y$ is the fraction of bound component being monitored via fluorescence signal.

#### 1.9 Fluorescence quenching experiments

Fluorescence quenching experiments were performed following the previously published procedure[3]. Mixtures were prepared as an equimolar ratio of acidic and basic peptide analogues at 50 µM of the 4-cyanophenylalanine (4CF) containing peptide. For both 4CF peptides, fluorescence was first tested in isolation before mixing with an equivalent concentration of MSe containing peptides. Samples were prepared at room temperature and were also annealed at 95 °C for 60 seconds, then slowly cooled back to room temperature over 2 hours. Fluorescence experiments were then conducted using a JASCO FP-6500 spectrofluorometer (Hachioji, Tokyo) and with either 26.50-

F/Q/10 or 26.160-F/Q/10 quartz cuvettes provided by Starna Scientific (Hainault, UK). Samples were excited at 240 nm with a bandwidth of 3 nm whilst readings were taken from 260 – 400 nm at a bandwidth of 1 nm reading 200 nm/min. A 1 s response time was used with a 0.5 nm data pitch. A manual voltage of 500 was used to amplify the signal. The experiments were conducted in 25 mM HEPES at pH 7.5 in the absence of NaCl as chloride ions are known to quench 4CF fluorescence.[3]

#### 1.10 Crystal growth

Diffraction-quality crystals were grown using a sitting-drop vapor-diffusion method. Freeze-dried peptides were resuspended in deionized water to concentrations of 10 mg ml$^{-1}$. CC-Hex2-A-3-***g*** and CC-Hex2-B-3-***g*** were mixed in a 1:1 ratio and annealed by heating to 95 ºC and slowly cooling back to room temperature. Commercially available sparse matrix screens were used (JCSG-plusTM, Structure Screen 1 + 2, ProPlexTM, Morpheus® and PACT PremierTM), and the drops were dispensed using a robot (Oryx8; Douglas Instruments). For each well of an MRC 2 drop plate, 0.3 µL of the peptide solution were equilibrated with 0.3 µL of the screen solution in parallel with 0.4 µL of the peptide solution equilibrated with 0.2 µL of the screen solution and the plate was incubated at 20 ºC. To aid with cryoprotection, crystals were soaked in their respective reservoir solutions containing 25% glycerol prior to freezing.

#### 1.11 X-ray crystal structure determination

X-ray diffraction data were collected at Diamond Light Source (Didcot, UK) on beamline I24 (Supplementary Table 2). Data were processed using the automatedXia2 pipeline, which ports data through DIALS (2.0.2) to POINTLESS (1.11.1) and AIMLESS (0.5.32) as implemented in the CCP4 suite. The structure was phased using *ab initio* phasing using ARCIMBOLDO_LITE. The initial phases were modelled into and refined using BUCCANEER. The final structure was obtained after iterative rounds of model building with COOT and refinement with REFMAC5 (7.1) and Phenix Refine (1.19.2_4158). TLS parameters were used during refinement as one group per chain for all structures. Torsion NCS restraints were used for fragments with <2Å RMSD and 90% sequence identity. Solvent-exposed atoms lacking map density were either deleted or left at full occupancy. Data collection and refinement statistics are provided in Supplementary Table 2.

#### 1.12 Multimer structure prediction

AlphaFold-Multimer (AF-M) predictions were generated using ColabFold (version 1.5.2) without MSAs.[4-6] 100 seeds for all 5 models were sampled, generating 500 predictions per sequence. Amber was used to minimize the top 5 ranking predictions on a NVIDIA RTX A5500. All other parameters for AF-M were kept at default values and are shown in the JSON configuration below:

```json
{
    "num_queries": 3,
    "use_templates": false,
    "num_relax": 5,
    "msa_mode": "single_sequence",
    "model_type": "alphafold2_multimer_v3",
    "num_models": 5,
    "num_recycles": null,
    "recycle_early_stop_tolerance": null,
    "num_ensemble": 1,
    "model_order": [1,2,3,4,5],
    "keep_existing_results": true,
    "rank_by": "multimer",
    "max_seq": 1,
    "max_extra_seq": 1,
    "pair_mode": "unpaired_paired",
    "host_url": "https://api.colabfold.com",
    "stop_at_score": 100,
    "random_seed": 0,
    "num_seeds": 100,
    "recompile_padding": 10,
    "commit": "05c0cb38d002180da3b58cdc53ea45a6b2a62d31",
    "use_dropout": false,
    "use_cluster_profile": true,
    "use_fuse": true,
    "use_bfloat16": true,
    "version": "1.5.2"
}
```

The CC-Hex2-AB-3-***g*** computed structure models were clustered based on structure similarity using template modelling scores (TMscores).[7] TMscores were subsequently normalized for fraction aligned (normalized TMscore = TMscore × Fraction aligned) to punish incomplete alignments. Results were binned into clusters with 0.95 Normalised TMscore similarity.

AlphaFold3 predictions were run using the public webserver. For analysis the predicted structure with the best `ranking_score` is used. All predictions were run with seed 42.

### 2 SUPPLEMENTARY DATA

Table S1. MALDI-TOF and analytical ultracentrifugation statistics of designed peptides.

| Name | Monomeric mass expected (g/mol) | Monomeric mass observed MALDI-TOF (g/mol) | Partial specific volume ($\bar{v}$, $cm^3$ $g^{-1}$) | Fitted mass AUC-SV (95 % confidence, 3SF, g/mol) | f/f0 | s (S) | s20,w (S) | Fitted mass AUC-SE (95 % confidence, 3SF, g/mol) | AUC-SE molecular mass / monomer mass |
|---|---|---|---|---|---|---|---|---|---|
| CC-Hex2-A-4-***c*** | 3314.7 | 3316.6 | | | | | | | |
| CC-Hex2-B-4-***c*** | 3307.2 | 3308.4 | | | | | | | |
| CC-Hex2-A-3-***a*** | 2543.8 | 2566.8 | 0.765 | 17,200, 34,200 | 1.544, 1.544 | 1.273, 2.014 | 1.333, 2.110 | | |
| CC-Hex2-B-3-***a*** | 2515.1 | 2516.0 | | | | | | | |
| CC-Hex2-A-3-***b*** | 2543.8 | 2582.3 | 0.776 | 11,800 | 1.148 | 1.278 | 1.322 | 14,300 | 5.7 |
| CC-Hex2-B-3-***b*** | 2515.1 | 2539.2 | | | | | | | |
| CC-Hex2-A-3-***b***-MSe | 2576.7 | 2600.4 | | | | | | | |
| CC-Hex2-B-3-***b***-CNF1 | 2523.9 | 2547.1 | | | | | | | |
| CC-Hex2-B-3-***b***-CNF2 | 2587.1 | 2604.3 | | | | | | | |
| CC-Hex2-A-3-***c*** | 2543.8 | 2582.0 | 0.765 | 11,200 | 1.173 | 1.256 | 1.316 | 15,400 | 6.1 |
| CC-Hex2-B-3-***c*** | 2515.1 | 2537.3 | | | | | | | |
| CC-Hex2-A-3-***c***-MSe | 2576.8 | 2601.2 | | | | | | | |
| CC-Hex2-B-3-***c***-CNF1 | 2523.9 | 2524.7 | | | | | | | |
| CC-Hex2-B-3-***c***-CNF2 | 2524.0 | 2524.7 | | | | | | | |
| CC-Hex2-A-3-***d*** | 2543.8 | 2567.2 | 0.765 | 13,000, 31,033 | 1.103, 1.103 | 1.479, 2.640 | 1.549, 2.766 | | |
| CC-Hex2-B-3-***d*** | 2515.1 | 2516.0 | | | | | | | |
| CC-Hex2-A-3-***e*** | 2543.8 | 2582.0 | | | | | | | |
| CC-Hex2-B-3-***e*** | 2515.1 | 2537.7 | | | | | | | |
| CC-Hex2-A-3-***f*** | 2543.8 | 2566.9 | | | | | | | |
| CC-Hex2-B-3-***f*** | 2515.1 | 2515.2 | | | | | | | |
| CC-Hex2-A-3-***g*** | 2543.8 | 2517.2 | 0.76 | 14,200 | 1.200 | 1.482 | 1.548 | 15,200 | 6.0 |
| CC-Hex2-B-3-***g*** | 2515.1 | 2568.5 | | | | | | | |
| CC-Hex2-A-3-***g***-MSe | 2576.8 | 2600.9 | | | | | | | |
| CC-Hex2-B-3-***g***-CNF1 | 2523.9 | 2524.7 | | | | | | | |
| CC-Hex2-B-3-***g***-CNF2 | 2524.0 | 2548.1 | | | | | | | |

Table S2. Crystal structure refinement statistics for CC-Hex2-AB-3-*g*.

| | CC-Hex2-AB-3-*g* |
|---|---|
| **PDB ID** | 9EVG |
| **Data Collection** | |
| Source | Diamond I24 |
| Detector | PILATUS3 6M |
| Wavelength | 0.98 |
| Resolution range | 47.31 - 1.903 (1.971 - 1.903) |
| Space group | P 1 21 1 |
| Unit cell: *a, b, c (Å)* | 32.6756 94.6118 51.1156 |
| *α, β, γ (°)* | 90 91.7522 90 |
| Total reflections | 321327 (31423) |
| Unique reflections | 24363 (2402) |
| Multiplicity | 13.2 (13.1) |
| Completeness (%) | 99.86 (99.26) |
| Mean I/sigma(I) | 11.57 (1.59) |
| Wilson B-factor | 29.29 |
| R-merge | 0.114 (0.2568) |
| R-meas | 0.1189 (0.2673) |
| R-pim | 0.03329 (0.0735) |
| CC1/2 | 0.997 (0.975) |
| CC* | 0.999 (0.994) |
| **Refinement** | |
| Reflections used in refinement | 24349 (2402) |
| Reflections used for R-free | 1134 (95) |
| R-work | 0.1921 (0.2743) |
| R-free | 0.2185 (0.3311) |
| CC(work) | 0.956 (0.809) |
| CC(free) | 0.937 (0.645) |
| Number of non-hydrogen atoms | 2020 |
| macromolecules | 1796 |
| ligands | 121 |
| solvent | 147 |
| Protein residues | 272 |

| | |
|---|---|
| RMS(bonds) | 0.004 |
| RMS(angles) | 0.45 |
| Ramachandran favored (%) | 99.6 |
| Ramachandran allowed (%) | 0.4 |
| Ramachandran outliers (%) | 0 |
| Rotamer outliers (%) | 0 |
| Clashscore | 1.09 |
| Average B-factor | 72.69 |
| macromolecules | 71.58 |
| ligands | 93.15 |
| solvent | 75.63 |
| Number of TLS groups | 12 |

#### 2.1 MALDI-TOF and analytical HPLC

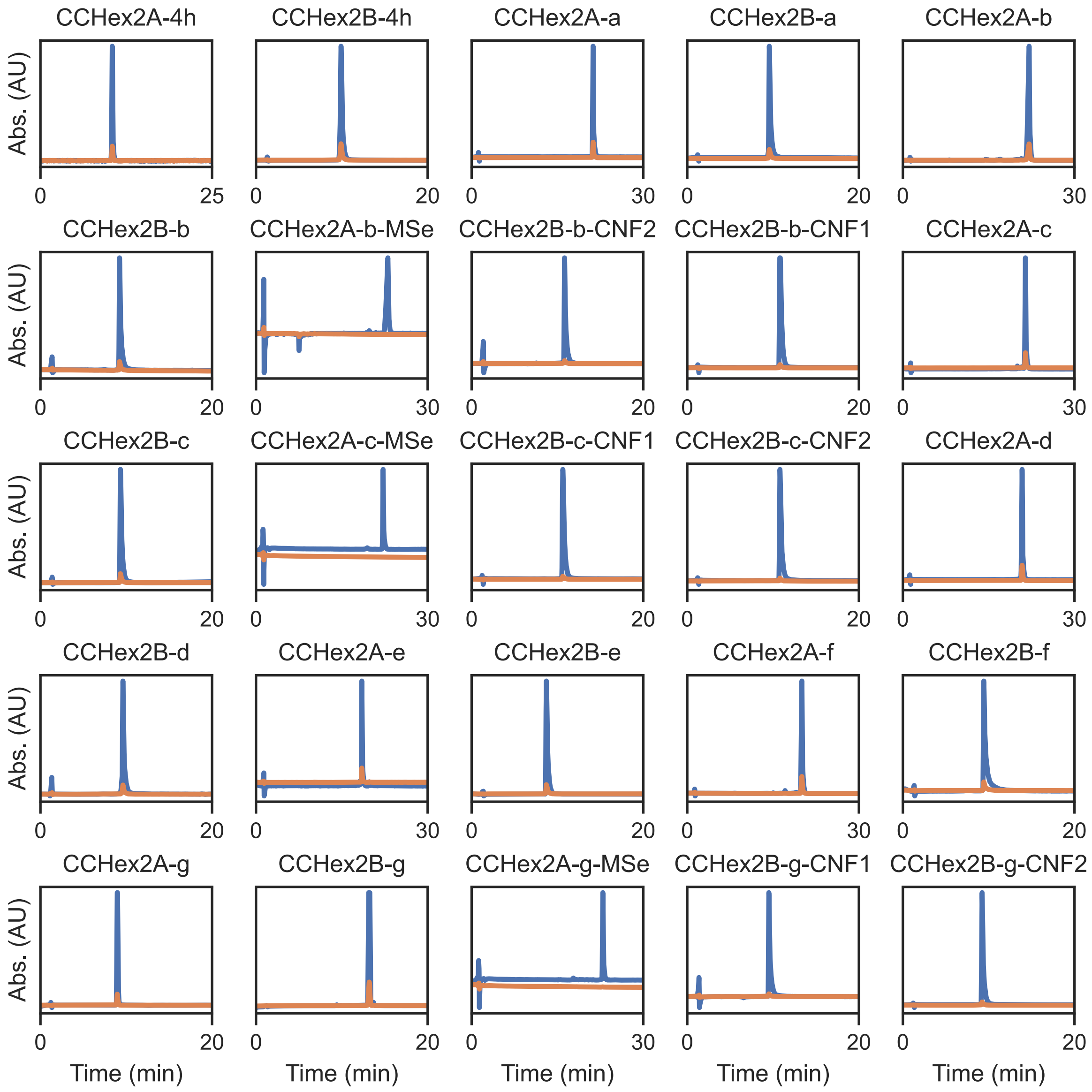

Figure S1. HPLC traces (blue 220, orange 280 nm) of peptides designed for this study. All traces are corrected for baseline absorbance.

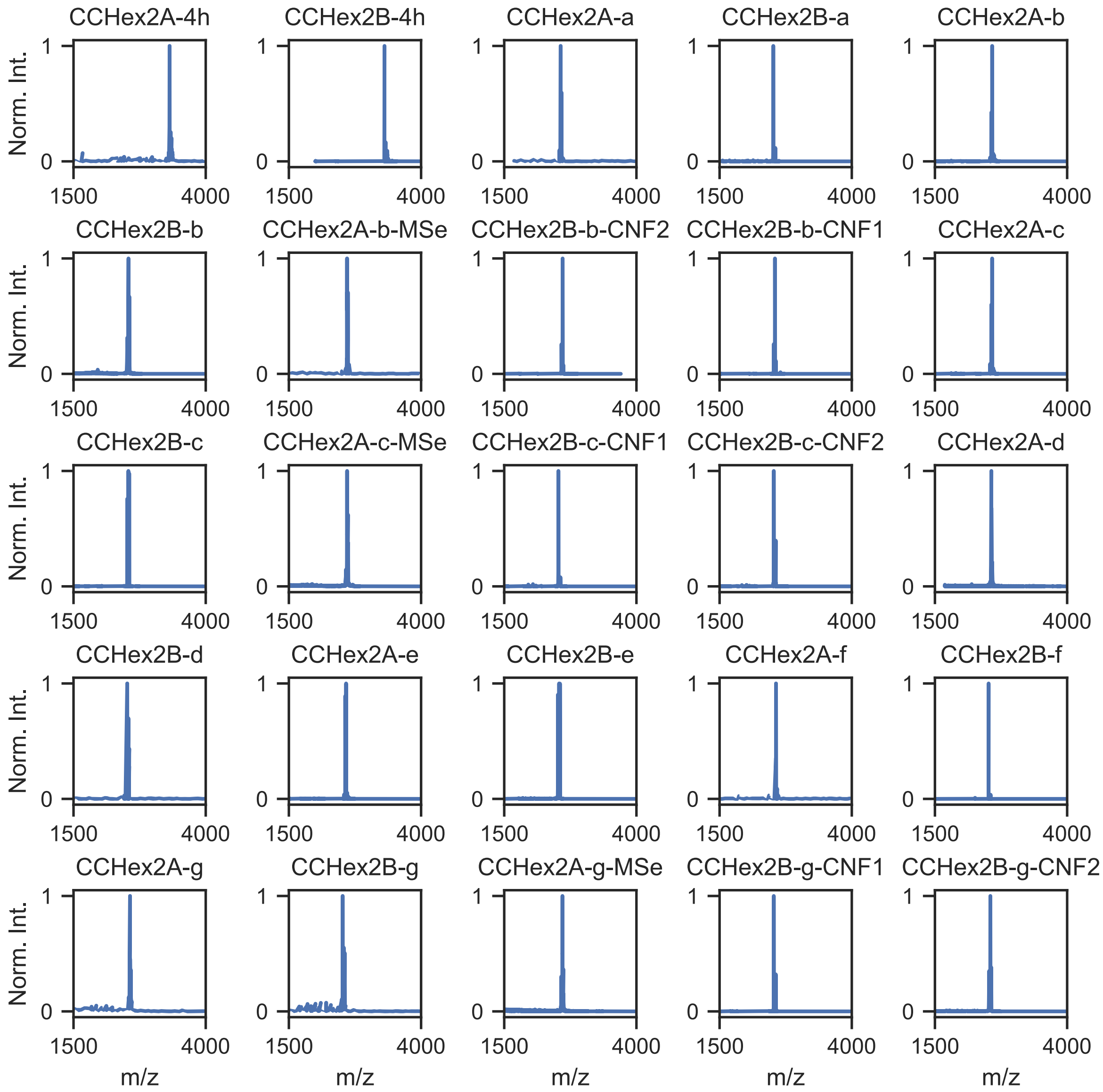

Figure S2. MADLI-TOF MS of peptides designed for this study. Observed masses are shown in Supplementary table 1.

#### 2.2 Circular Dichroism (CD) spectroscopy

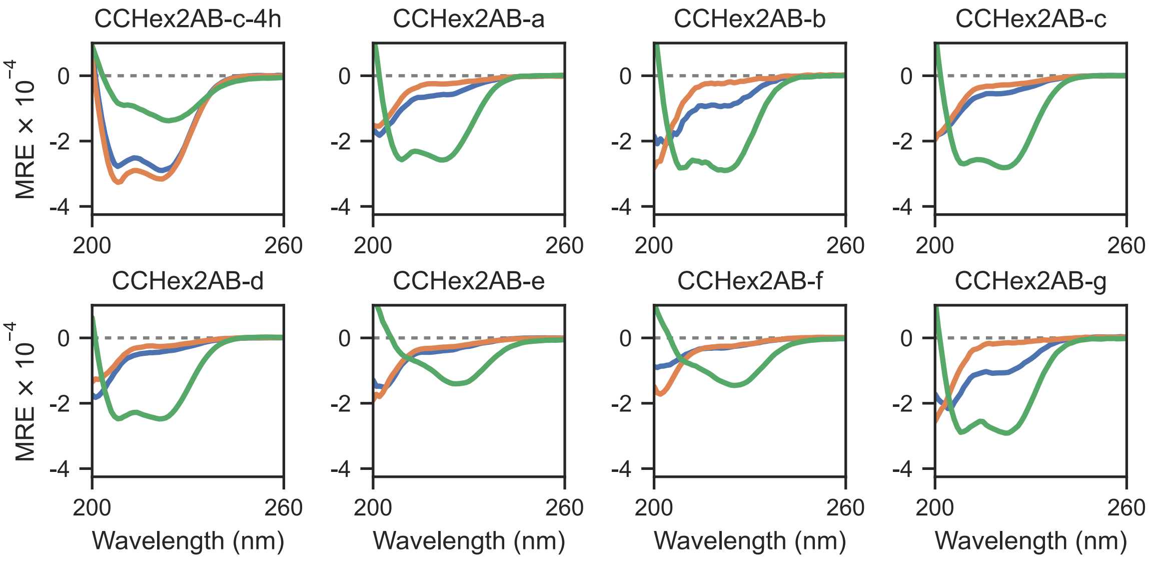

Figure S3. CD spectra at 20 °C for peptides designed for this study. Acidic peptides are shown in blue, basic in orange and mixtures in green. MRE, mean residue ellipticity (deg cm2 dmol$^{-1}$ res$^{-1}$). Conditions: 100 µM peptide (50:50 when mixed), HBS.

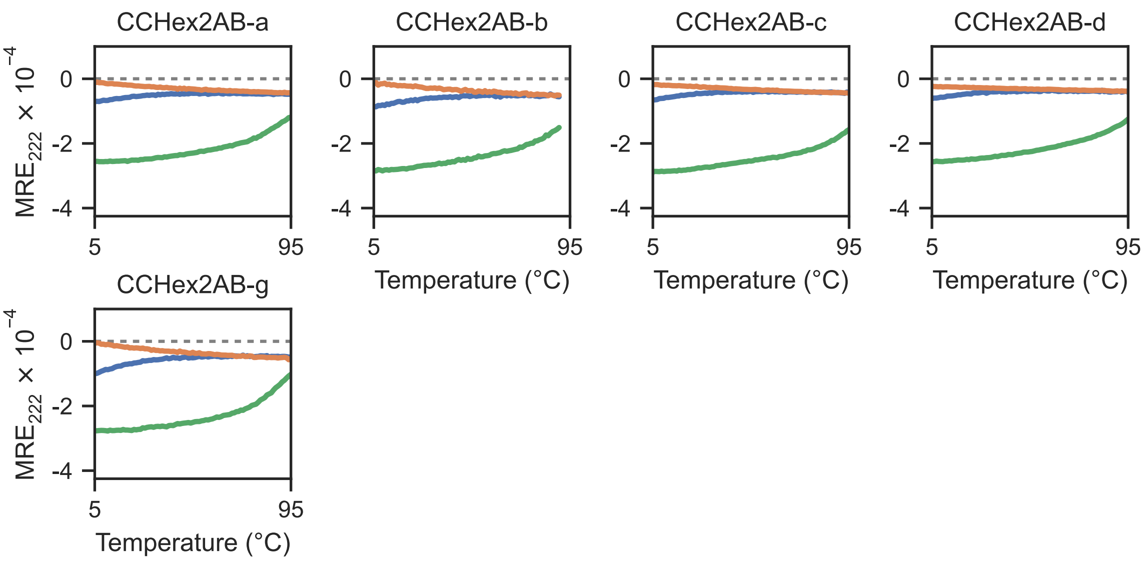

Figure S4. Temperature dependent CD signal, monitored at 222 nm for helical peptides designed for this study. Acidic peptides are shown in blue, basic in orange and mixtures in green. MRE, mean residue ellipticity (deg cm2 dmol$^{-1}$ res$^{-1}$). Conditions: 100 µM peptide (50:50 when mixed), HBS.

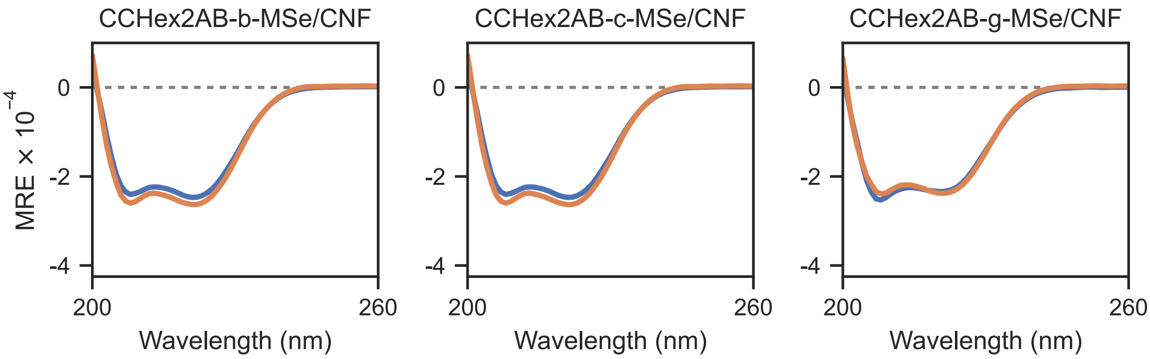

Figure S5. CD spectra at 20 °C for MSe and CNF containing peptides designed for this study. Both combinations of CNF variants are shown; CCHex2A-***x***-MSe + CCHex2B-***x***-CNF1 (blue), CCHex2A-***x***-MSe + CCHex2B-***x***-CNF2 (orange), ***x*** denoting helix register (***b***/***c***/***g***). MRE, mean residue ellipticity (deg cm2 dmol$^{-1}$ res$^{-1}$). Conditions: 100 µM peptide (50:50 when mixed), HBS.

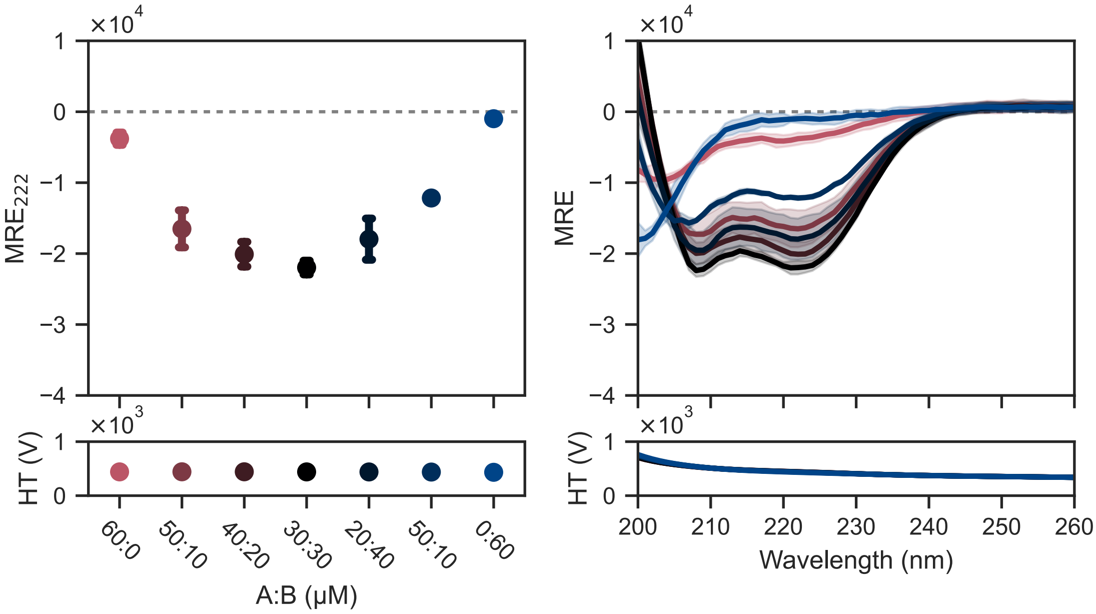

Figure S6. Concentration dependent CD signal monitored at 222 nm, (left) and CD spectra at 20 °C (right) for CCHex2AB-b. Color denotes the ratio of CCHex2A-b:CCHex2B-b (red 1:0, black 1:1, blue 0:1). MRE, mean residue ellipticity (deg cm2 dmol-1 res-1). Conditions: HBS. Markers show the mean of the data and range bars represent one standard deviation of the mean. *N* = 3.

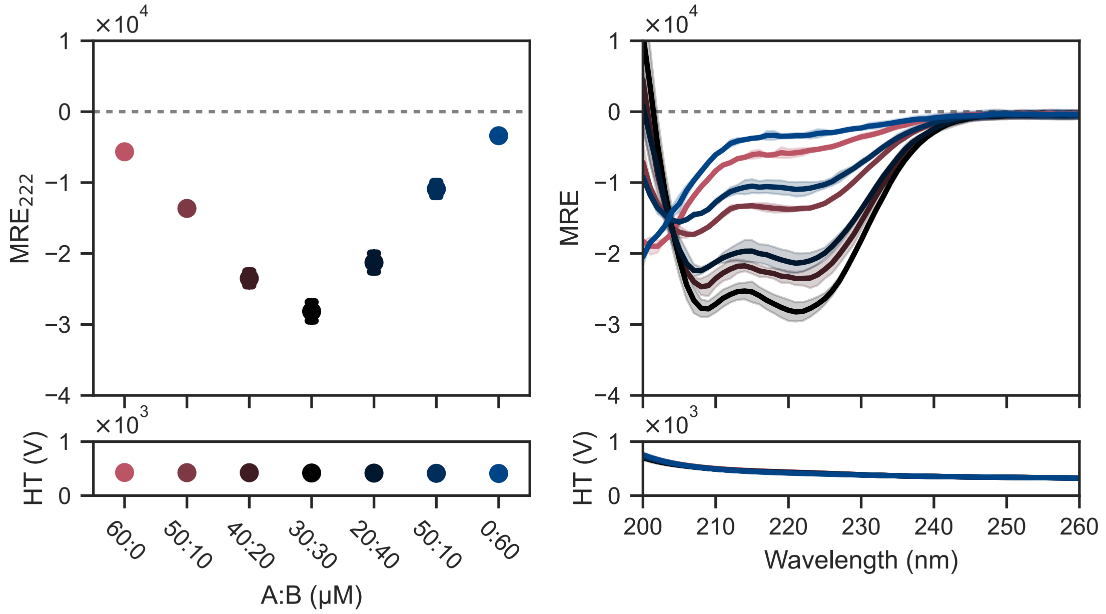

Figure S7. Concentration dependent CD signal monitored at 222 nm, (left) and CD spectra at 20 °C (right) for CCHex2AB-c. Color denotes the ratio of CCHex2A-c:CCHex2B-c (red 1:0, black 1:1, blue 0:1). MRE, mean residue ellipticity (deg cm2 dmol-1 res-1). Conditions: HBS. Markers show the mean of the data and range bars represent one standard deviation of the mean. N = 2.

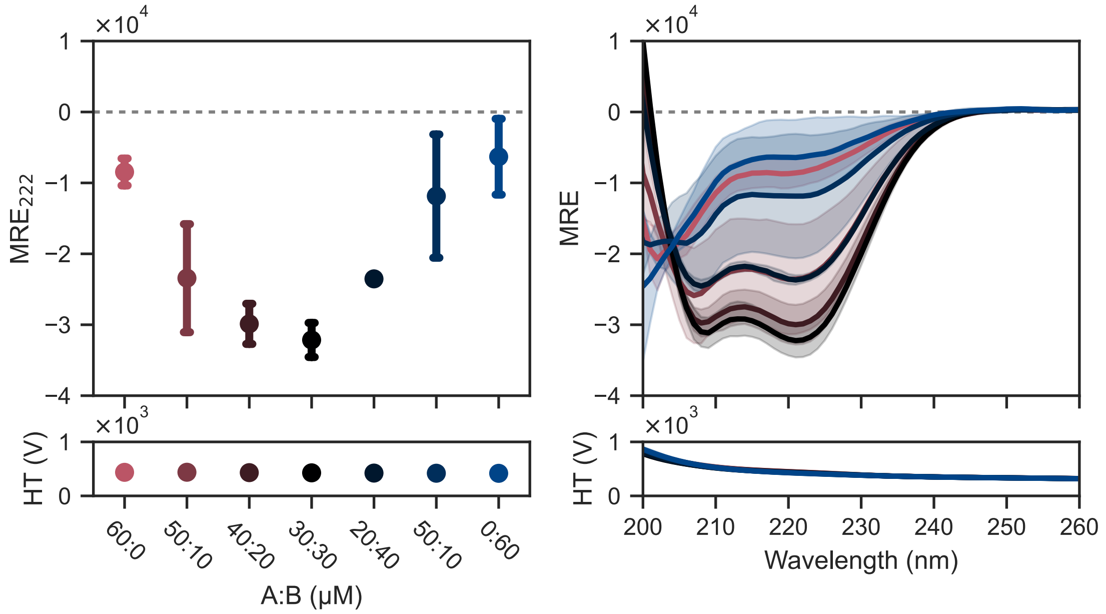

Figure S8. Concentration dependent CD signal monitored at 222 nm, (left) and CD spectra at 20 °C (right) for CCHex2AB-g. Color denotes the ratio of CCHex2A-g:CCHex2B-g (red 1:0, black 1:1, blue 0:1). MRE, mean residue ellipticity (deg cm2 dmol-1 res-1). Conditions: HBS. Markers show the mean of the data and range bars represent one standard deviation of the mean. N = 3.

#### 2.3 Analytical Ultracentrifugation (AUC)

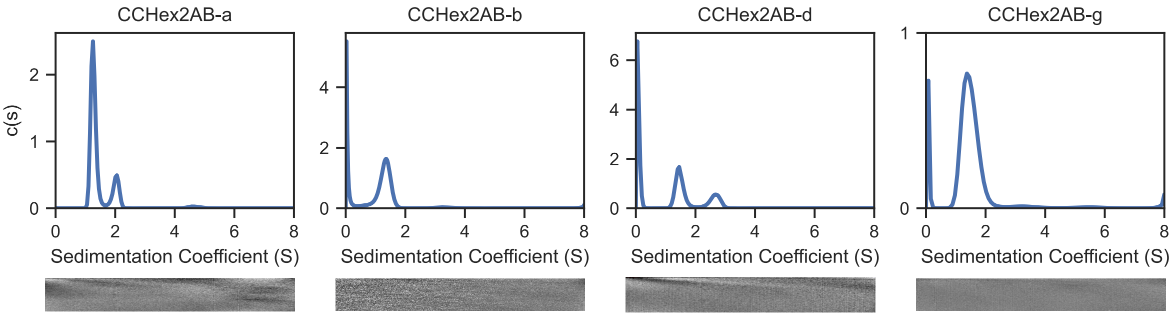

Figure S9. Analytical Ultracentrifugation sedimentation velocity traces of peptides designed for this study. Residuals are shown as a bitmap (below). Results for individual peptides can be found in Supplementary table 1. Conditions: 75 μM CCHex2A-***x*** + 75 μM CCHex2B-***x***, HBS.

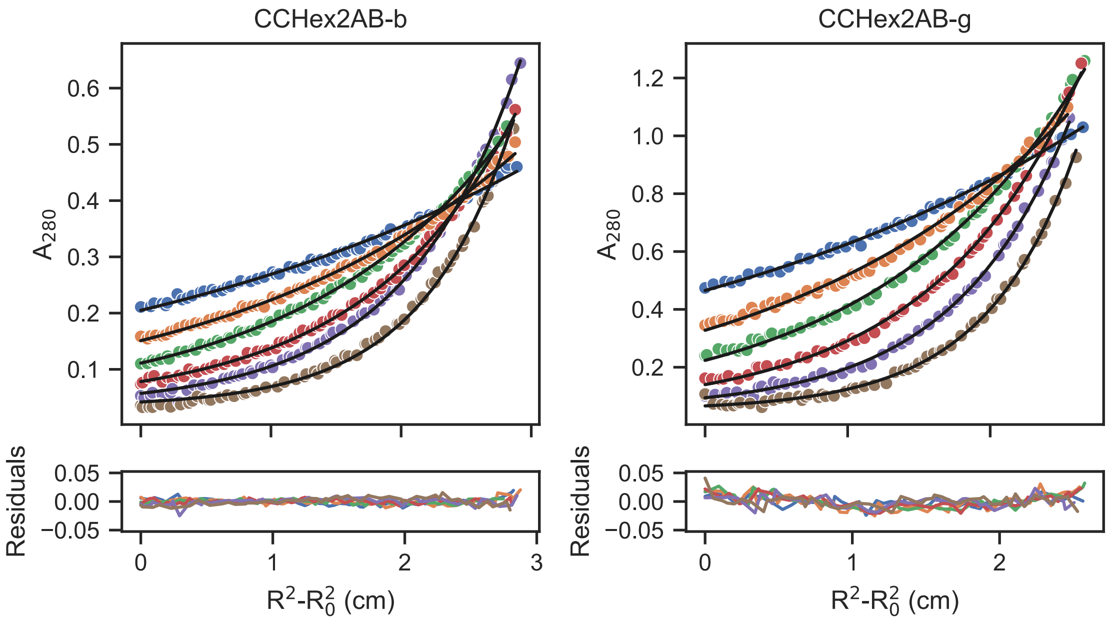

Figure S10. Analytical Ultracentrifugation sedimentation equilibrium traces of peptides designed for this study. Data (markers) and fit (solid lines) are shown in the upper plots with residuals shown below. Colors indicate rotor speed (rpm): 20,000 (blue), 25,000 (orange), 30,000 (green), 35,000 (red), 40,000 (purple), 45,000 (brown). Results for individual peptides can be found in Supplementary table 1. Conditions: 35 μM CCHex2A-***x*** + 35 μM CCHex2B-***x***, HBS.

#### 2.4 Ligand Binding assays

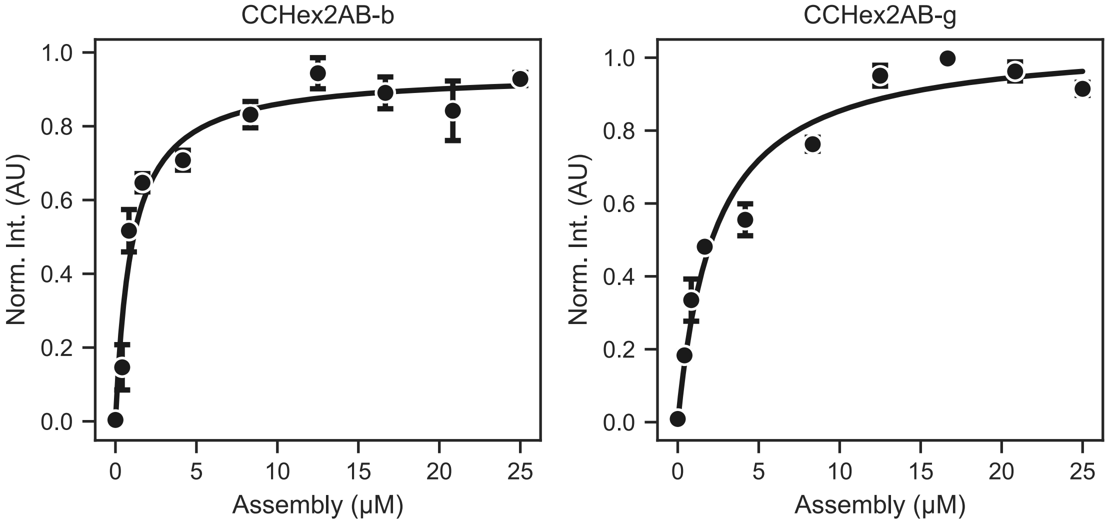

Figure S11. Saturation binding curves with DPH for tested peptides. Plots show mean fluorescence data (markers) with standard deviations (error bars). N = 3. Curves represent fits to a single site binding model. Peptide concentration is converted to αHB assembly concentration by the oligomeric state. CCHex2AB-b results: KD = 6.0 μM (± 1.7), R2 = 0.959. CCHex2AB-g: KD = 13.9 μM (± 8.7), R2 = 0.969. Conditions: 2.5 – 150.0 μM peptide concentrations (equimolar), HBS, 1 μM DPH, 5 % v/v DMSO.

#### 2.5 AlphaFold3 computed structure models

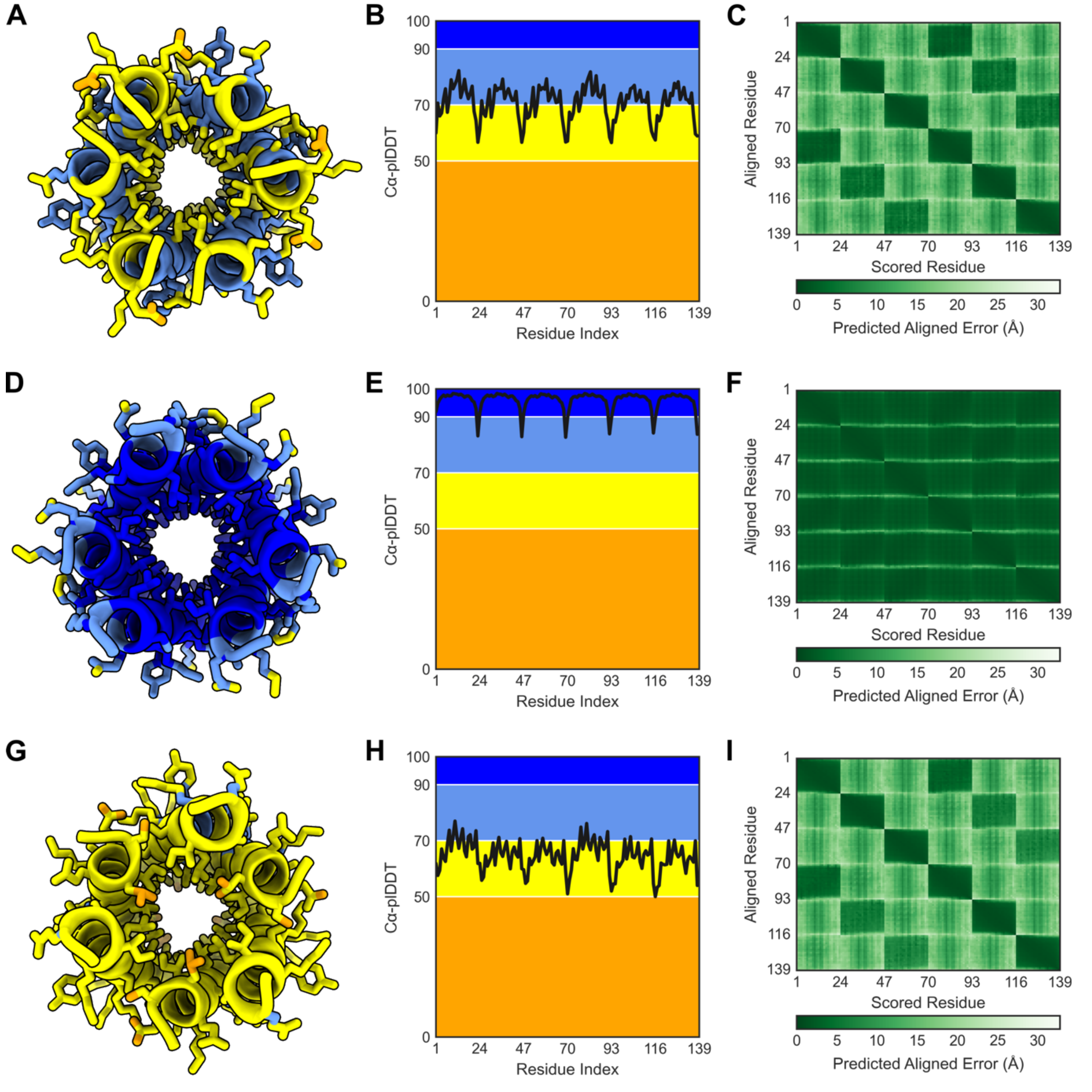

Figure S12. AlphaFold3 results for CCHex2-AB-***b*** (A – C), CCHex2-AB-***c*** (D – F) and CCHex2-AB-***g*** (G – I). (A,D,G) All-atom prediction colored by atom plDDT confidence (very low: plDDT < 50, orange; low: 50 < plDDT < 70, yellow; confident: 70 < plDDT < 90, light blue; very high: 90 < plDDT, blue). (B,E,H) The Cα-plDDT scores for each residue. Patches reflect confidence bins as described for A,D,G. (C,F,I) The predicted aligned error (pAE) at each residue index.

### 3 REFERENCES

(1) Kuipers, B. J. H.; Gruppen, H. (2007) Prediction of Molar Extinction Coefficients of Proteins and Peptides Using UV Absorption of the Constituent Amino Acids at 214 nm To Enable Quantitative Reverse Phase High-Performance Liquid Chromatography−Mass Spectrometry Analysis. *J. Agric. Food Chem. 55,* 5445-5451.

(2) http://www.analyticalultracentrifugation.com/sedphat/sedphat.html (accessed 2019 October).

(3) Watson, M. D.; Peran, I.; Raleigh, D. P. (2016) A Non-perturbing Probe of Coiled Coil Formation Based on Electron Transfer Mediated Fluorescence Quenching. *Biochemistry 55,* 3685-3691.

(4) Jumper, J.; Evans, R.; Pritzel, A.; Green, T.; Figurnov, M.; Ronneberger, O.; Tunyasuvunakool, K.; Bates, R.; Žídek, A.; Potapenko, A.; et al. (2021) Highly accurate protein structure prediction with AlphaFold. *Nature 596,* 583-589.

(5) Mirdita, M.; Schütze, K.; Moriwaki, Y.; Heo, L.; Ovchinnikov, S.; Steinegger, M. (2022) ColabFold: making protein folding accessible to all. *Nat. Methods* 1-4.

(6) Evans, R.; O'Neill, M.; Pritzel, A.; Antropova, N.; Senior, A.; Green, T.; Žídek, A.; Bates, R.; Blackwell, S.; Yim, J.; et al. (2022) Protein complex prediction with AlphaFold-Multimer. *bioRxiv* 2021.2010.2004.463034.

(7) Zhang, Y.; Skolnick, J. (2004) Scoring function for automated assessment of protein structure template quality. *Proteins 57,* 702-710.
